# Disruption of *miR-18a* alters proliferation, photoreceptor replacement kinetics, inflammatory signaling and microglia/macrophage numbers during retinal regeneration in zebrafish

**DOI:** 10.1101/2021.04.25.441353

**Authors:** Evin Magner, Pamela Sandoval-Sanchez, Ashley C. Kramer, Ryan Thummel, Peter F. Hitchcock, Scott M. Taylor

**Author notes:** Correspondence to Scott M. Taylor.

## Abstract

In mammals, photoreceptor loss causes permanent blindness, but in zebrafish (*Danio rerio*), photoreceptor loss reprograms Müller glia to function as stem cells, producing progenitors that regenerate photoreceptors. MicroRNAs (miRNAs) regulate CNS neurogenesis, but the roles of miRNAs in injury-induced neuronal regeneration are largely unknown. In the embryonic zebrafish retina, *miR-18a* regulates photoreceptor differentiation. The purpose of the current study was to determine, in zebrafish, the function of *miR-18a* during injury-induced photoreceptor regeneration. RT-qPCR, in-situ hybridization and immunohistochemistry showed that *miR-18a* expression increases throughout the retina between 1 and 5 days post-injury (dpi). To test *miR-18a* function during photoreceptor regeneration, we used homozygous *miR-18a* mutants (*miR-18a^mi5012^*), and knocked down *miR-18a* with morpholino oligonucleotides. During photoreceptor regeneration, *miR-18a^mi5012^* retinas have fewer mature photoreceptors than WT at 7 and 10 dpi, but there is no difference at 14 dpi, indicating that photoreceptor regeneration is delayed. Labeling dividing cells with 5-bromo-2′-deoxyuridine (BrdU) showed that at 7 and 10 dpi, there are excess dividing progenitors in both mutants and morphants, indicating that *miR-18a* negatively regulates injury-induced proliferation. Tracing 5-ethynyl-2′-deoxyuridine (EdU) and BrdU-labeled cells showed that in *miR-18a^mi5012^* retinas excess progenitors migrate to other retinal layers in addition to the photoreceptor layer. Inflammation is critical for photoreceptor regeneration, and RT-qPCR showed that in *miR-18a^mi5012^* retinas, inflammatory gene expression and microglia activation are prolonged. Suppressing inflammation with dexamethasone rescues the *miR-18a^mi5012^* phenotype. Together, these data show that in the injured zebrafish retina, disruption of *miR-18a* alters proliferation, inflammation, the microglia/macrophage response, and the timing of photoreceptor regeneration.

## INTRODUCTION

In humans, photoreceptor death causes permanent blindness. In contrast, in zebrafish, *Danio rerio*, photoreceptor death stimulates Müller glia (MG) to reprogram into a stem cell state and divide, thereby producing neuronal progenitors that fully regenerate the lost photoreceptors [1–4]. This regenerative capacity has established the zebrafish retina as an outstanding model for investigating photoreceptor degeneration and regeneration [5]. Mechanisms identified in the zebrafish retina that govern the reprogramming of Müller glia and neuronal regeneration have been used to develop methods to stimulate some neurons to regenerate in the mouse retina [6–8]. Understanding the mechanisms that govern photoreceptor regeneration in zebrafish could, therefore, be critical for developing regenerative therapies to treat human blindness.

Recent research has improved our understanding of the molecular pathways that regulate neuronal regeneration in the zebrafish retina [reviewed in 4,3,9,10]. The primary focus of this research has been on identifying the transcriptional mechanisms involved in neuronal regeneration, but recent studies show that post-transcriptional regulation by non-coding RNAs, and specifically microRNAs (miRNAs), also play critical roles in retinal neuronal regeneration [11–15]. Of the more than 2600 mature miRNAs coded by the vertebrate genome [16], the functional roles of a very small number have been established in retinal regeneration. MicroRNAs are likely to play key roles in regulating photoreceptor regeneration, and additional studies that determine the roles of miRNAs in photoreceptor regeneration are critically needed.

Inflammation occurs in response to tissue injury, and several recent studies show that activation of neuroinflammatory pathways is both necessary and sufficient to initiate neuronal (including photoreceptor) regeneration in the zebrafish retina [17–22] [reviewed in 23,24]. Inflammatory signals seem to have the opposite effect in the injured mammalian retina, in which ablating microglia reduces inflammatory gene expression and increases the number of regenerated neurons [25]. Understanding mechanisms that control inflammation in the injured retina may, therefore, be critical to unlocking the regenerative potential in the mammalian retina. Importantly, several miRNAs have been identified as key regulators of inflammatory pathways [26] and as biomarkers of inflammatory disease [27], and identifying the roles of miRNAs in the injured retina could be critical for understanding the link between the retinal inflammatory response and neuronal regeneration.

MicroRNAs are small, 18-25 nucleotides long, non-coding RNAs that generally function by binding to the 3’ untranslated region of mRNA and inhibit translation and/or promote mRNA degradation [28, 29]. The miRNA, *miR-18a*, was recently found to regulate photoreceptor differentiation in larval zebrafish by suppressing levels of the transcription factor, NeuroD [30]. The role of *miR-18a* during photoreceptor regeneration in zebrafish is currently unknown. Bioinformatics tools predict that *miR-18a* can interact directly with mRNAs that encode more than 25 molecules that function in inflammatory pathways (http://www.targetscan.org/fish_62/), suggesting that *miR-18a* might be an important regulator of inflammation. However, *miR-18a* has not been investigated in the context of inflammatory regulation.

The objective of this research was to determine if *miR-18a* is necessary for normal photoreceptor regeneration and inflammatory pathway regulation in the injured zebrafish retina. qPCR showed that *miR-18a* expression increases between 3 and 5 days post injury (dpi) and then decreases by 7 dpi. *In situ* hybridization for *miR-18a*, combined with green fluorescent protein (GFP) immunolabeling in transgenic *Tg*(*gfap:egfp^mi2002^*) fish [31] with GFP-labeled Müller glia showed that the *miR-18a* expression increases throughout the retina and is expressed in Müller glia (MG) by 1 dpi and in MG-derived progenitors by 3 dpi. 5-Bromo-2’-deoxyuridine (BrdU) labeling revealed that relative to wild-type (WT) animals, at 7 and 10 dpi, homozygous *miR-18a* mutants (*miR-18a^mi5012^*) have significantly more proliferating MG-derived progenitors. The regeneration of rods and cones is initially delayed in *miR-18a^mi5012^* fish, but numerically matches wild type animals by 14dpi. There was no overproduction of regenerated photoreceptors. BrdU and 5-ethynyl-2’-deoxyuridine (EdU) tracing showed that in *miR-18a^mi5012^* retinas compared with WT, a lower proportion of Müller glia-derived progenitors differentiate into photoreceptors and more migrate to various layers of the retina, in addition to the outer nuclear layer (ONL), suggesting that the excess progenitors may differentiate into other cell types besides photoreceptors. RT-qPCR and *in situ* hybridization showed that at 5 and 7 dpi, when inflammation is normally resolving, in *miR-18a^mi5012^* retinas, the expression of genes encoding pro-inflammatory cytokines and the cytokine regulator, *nfkb1*, are significantly elevated. Finally, suppressing inflammation with dexamethasone in *miR-18a^mi5012^* fish fully rescues both the excess proliferation and the delay in the regeneration of cone photoreceptors. Together, these data show that following photoreceptor injury in zebrafish, loss of *miR-18a* results in excess proliferation of progenitor cells, delayed photoreceptor regeneration, and a prolonged microglia/macrophage and inflammatory response. These results suggest a functional role for *miR-18a* in regulating injury-induced photoreceptor regeneration by regulating the resolution phase of inflammation.

## METHODS

### Fish husbandry, photolytic lesions and tissue preparation

All fish were maintained at 28.5°C on a 14/10-h light/dark cycle under standard husbandry conditions [32]. AB wild-type (WT) strain zebrafish, purchased from the Zebrafish International Research Center (ZIRC; University of Oregon, Portland, OR, USA), were used for control experiments. The *miR-18a^mi5012^* line has a 25 bp insertion in the sequence coding for the precursor molecule *pre-miR-18a*, and homozygous mutant fish, used for all experiments here, do not produce mature *miR-18a* [30]. The transgenic line, *Tg(gfap:egfp)^mi2002^* [31], expresses Enhanced Green Fluorescent Protein (EGFP) in Müller glia and MG-derived progenitors, and was used in conjunction with *in situ* hybridization to determine the cellular expression of *miR-18a*.

As previously described, photolytic lesions were used to selectively kill photoreceptors [33]. Briefly, fish were exposed to ultra-high intensity light (>120,000 lux) in a 100 ml beaker for 30 minutes, using a SOLA SE II 365 White Light Engine (Lumencor, Beaverton, OR, USA). Following photolytic lesions, fish were maintained in normal system water and exposed to the standard 14/10-h light cycle.

Prior to collecting tissues, fish were euthanized in 0.1% Tricaine Methanesulfonate (MS-222) and then decapitated. For histology, eyes were removed, fixed (overnight at 4°C) in 4% paraformaldehyde, infiltrated with 20% sucrose (in PBS) and then in a 2:1 mixture of 20% sucrose and optimal cutting temperature (OCT) compound, and then finally embedded and frozen in OCT. Sections, 10μm in thickness, were collected through the center of the eye and thaw mounted onto glass slides. For qPCR, retinas were isolated from two fish per biological replicate (4 retinas), and total RNA, including small RNAs, was purified using the miRvana miRNA purification kit (AM1560; Invitrogen, Carlsbad, CA, USA).

### Systemic labeling with BrdU or EdU, dexamethasone treatment, immunohistochemistry and *in situ* hybridization

Cells in the S-phase of the cell cycle were labeled with BrdU by swimming fish in 5 mM BrdU for 24 or 48 hours, or intraperitoneal (IP) injection of EdU–10 μL injection of 1 mg/L EdU per fish in phosphate buffered saline (PBS), injections at 3 and 7 dpi. Fish were then either sacrificed immediately or at variable timepoints thereafter. For BrdU (and PCNA) immunolabeling, sections were incubated in 100°C sodium citrate buffer (10 mM sodium citrate, 0.05% Tween 20, pH 6.0) for 30 minutes to denature DNA and cooled at room temperature for 20 minutes. For EdU, labeling was visualized using the Click-it EdU kit (Invitrogen). Sections then were subjected to standard immunolabeling as described below.

Fish treated with dexamethasone to inhibit inflammation were exposed in system water to 15 mg/L dexamethasone (D1756; Sigma-Aldrich, Burlington, MA, USA) diluted in 0.1% MetOH [21]. Control fish were treated with 0.1% MetOH only. Fish were treated between 2-6 dpi. All solutions were changed daily, and fish were fed brine shrimp every other day.

Standard immunolabeling was performed using previously published protocols [34]. The primary and secondary antibodies and dilutions used here were: mouse anti-BrdU 1:100 (347580; BD Biosciences, Franklin Lakes, NJ, USA), 4c4 1:200 (mouse anti-fish leukocytes; 92092321; Sigma-Aldrich), mouse anti-PCNA 1:100 (MAB424MI; Sigma-Aldrich), rabbit anti-GFP (ab290; Abcam, Cambridge, UK), Zpr-1, 1:200 (anti-Arrestin 3, red-green double cones, ZIRC), Zpr-3, 1:200 (anti-Rhodopsin, rod photoreceptors, ZIRC), goat anti-mouse Alexa Fluor 555 1:500 (Invitrogen), and goat anti-rabbit Alexa Fluor 488 (Invitrogen). Following immunohistochemistry, when applicable, TUNEL labeling was performed using the Click-iT Plus TUNEL assay (C10617, Invitrogen).

*In situ* hybridizations were performed using previously published protocols [35, 36]. Riboprobes were generated from PCR products using the following primers and by adding a T3 polymerase sequence on the reverse primer (lowercase letters) [37]. Probes were generated for rods *rhodopsin* (F—GAGGGACCGGCATTCTACGTG, R— aattaaccctcactaaagggCTTCGAAGGGGTTCTTGCCGC) and cones *arr3a* (F—GAAGACCAGTGGAAATGGCCAG, R—aattaaccctcactaaagggTCAGAGGCAGCTCTACTGTCAC). *In situ* hybridization for mature *miR-18a* was performed using a miRCURY LNA detection probe (Exiqon/Qiagen, Germantown, MD), labeled with DIG at the 5′ and 3′ends. Standard *in situ* hybridization methods were used for *miR-18a*, as described above, but using a 0.25 μM probe working concentration at a hybridization temperature of 58°C. For comparisons of relative expression across post-injury time points or between WT and *miR-18a^mi5012^* fish, all tissue sections were placed on the same slides and/or developed for identical periods of time.

### Reverse transcriptase quantitative real-time PCR (RT-qPCR)

As described above, for RT-qPCR, retinas were isolated and total RNA purified from the retinas of 2 fish (4 retinas) per biological replicate. Three to five biological replicates were collected for each experimental group, treatment or time point. Reverse transcription was performed using the Superscript III First Strand cDNA Synthesis System (18080051, Invitrogen). Each biological replicate was run in triplicate using 20 ng cDNA and the Applied Biosystems PowerUp Sybr Green Master Mix (A25741, Applied Biosystems, Invitrogen), on an Applied Biosystems StepOnePlus 96-well Real-Time PCR System. The ΔΔCT method was used to calculate expression relative to WT (unlesioned or control) and data were normalized to *gpia* as the housekeeping gene. Sequences for the standard qPCR primers used were as follows: *gpia* F— TCCAAGGAAACAAGCCAAGC, R—TTCCACATCACACCCTGCAC; *nfkb1* F—CAGCTGGTGACCAACTCTCAG, R—TCCTGTAGGCCTCCATCATGC; *tnfα* F— CTGGAGAGATGACCAGGACCAGGCC, R—GCTGTGGTCGTGTCTGTGCCCAGTC; *il1β* F—GCATGAGGGCATCAGGCTGGAGATG, R— TCCGGCTCTCAGTGTGACGGCCTGC; *il6* F—CCTGTCTGCTACACTGGCTAC, R— CACTTCTGCCGGTCGCCAAGG; *pri-miR-18a* F— GGCTTTGTGCTAAGGTGCATCTAG, R—CAGAAGGAGCACTTAGGGCAGTAG. To quantify mature *miR-18a* expression, a TaqMan custom qPCR assay was designed for mature *miR-18a* and for the small nuclear RNA U6, to be used as the housekeeping gene for data normalization (ThermoFisher Scientific, Halethorp, MD, USA). The ΔΔCT method was used to calculate expression at different post-injury time points relative to unlesioned. Statistical significance between time points and between WT and *miR-18a^mi5012^* values for each gene was determined using a Student’s t-test (p<0.05).

### Morpholino injection and electroporation

For *miR-18a* RNA interference experiments, two lissamine-tagged morpholinos were utilized (Gene Tools LLC, Philomath, OR). Morpholinos were resuspended in nuclease-free water to a working concentration of 3 mM. An anti-*miR-18a* morpholino was used to target *miR-18a* (5′ –CTATCTGCACTAGATGCACCTTAG– 3′). This morpholino was previously confirmed to effectively knock down *miR-18a* in zebrafish embryos [38, 30]. The Gene Tools Standard Control morpholino (5′ – CCTCTTACCTCAGTTACAATTTATA – 3′) was used as a non-specific control, because it has no known target in the zebrafish genome. Morpholino injection and electroporation were performed as previously described [39]. Briefly, at 48 hrs post-light lesion, adult wild-type zebrafish were anesthetized, the outer most component of the cornea was removed with small forceps, a small incision was made in the peripheral cornea with a scalpel (Cat. #378235; Beaver-Visitec International, Inc., Waltham, MA) and 0.5 μl of a 3 mM morpholino solution was injected into the vitreous of the left eye using a Hamilton syringe outfitted with a 1.5 in, 33 g blunt-tipped needle (Cat #87930 and #7762-06; Hamilton Company, Reno, NV). Immediately following the injections, the left eye was electroporated with a CUY21 Square Wave Electroporator (Protech International, Inc., San Antonio, TX), using 2 consecutive 50msec pulses at 75V with a 1sec pause between pulses. A 3mm diameter platinum plate electrode (CUY 650-P3 Tweezers, Protech International, Inc.) was used to drive the morpholino into the central-dorsal region of the retina. Animals were sacrificed at 7 days post-light lesion and enucleated eyes were processed for immunohistochemistry.

### Cell counts and data analysis

All counts of BrdU-labeled progenitors and rod and cone photoreceptors were performed at 200x magnification across 0.3 mm of linear retina. Measurements and cell counts were done using ImageJ analysis software [40]. For each biological replicate (each fish), cells were counted in and averaged among 3 non-adjacent retinal cross-sections in the vicinity of the optic nerve. For each analysis, 3-5 biological replicates were used per treatment, time point or genotype. All comparisons were pairwise (e.g. WT vs. mutant) and Student’s t-tests were used to determine statistical significance (p values less than 0.05 were considered statistically signficant).

## RESULTS

### Following photoreceptor injury, *miR-18a* is expressed in both the inner and outer nuclear layers, including in Müller glia and Müller glia-derived progenitors

To initially determine if *miR-18a* might play a role in photoreceptor regeneration, RT-qPCR and *in situ* hybridization were performed to determine the temporal and spatial expression of *miR-18a* following photoreceptor injury. RT-qPCR quantification of the miRNA precursor, *pre-miR-18a*, showed that, compared with control uninjured retinas, *pre-miR-18a* expression is significantly higher at 3 and 5 days post-retinal injury (dpi) and Taqman qPCR showed that mature *miR-18a* expression is significantly higher at 3, 5, and 7 dpi (Figure 1a). *In situ* hybridization using an LNA ribroprobe for mature *miR-18a*, combined with immunolabeling for green fluorescent protein (GFP) in *Tg(gfap:egfp)^mi2002^* fish, showed that in control uninjured retinas, there is only very slight expression of *miR-18a* (Figure 1b) but by 24 hpi, around the time that Müller glia divide once to produce a neuronal progenitor [see 41], *miR-18a* is expressed in many cells throughout the inner nuclear layer (INL), including GFP-positive Müller glia (Figure 1c). Then at 3 dpi, during the peak of progenitor proliferation, *miR-18a* remains strongly expressed in the INL and ONL, including in GFP-positive Müller glia and MG-derived progenitors (Figure 1d). These progenitors are clearly visible as cells in the ONL that express GFP in their cytoplasm [described in 2], shown in Figure 1d with black outlined arrows. Finally, by 7 dpi, when MG-derived progenitors are normally exiting the cell cycle, *miR-18a* expression is again similar to the expression pattern seen between 0 and 24 hpi (Figure 1e, f). Focusing on the 3 dpi time point, at the peak of cell proliferation and *miR-18a* expression, in situ hybridization combined with proliferating cell nuclear antigen (PCNA) immunolabeling showed that *miR-18a* is expressed in GFP-labeled, PCNA-positive, proliferating MG-derived progenitors (Figure 2). At all time points, all GFP-labeled cells appear to express *miR-18a*, and the intensity of *miR-18a* labeling increases throughout the retina at 1, 3 and 5 dpi. Together, these results show that *miR-18a* expression is upregulated in the retina during the time periods when Müller glia and then MG-derived progenitors divide.

**Fig. 1.**
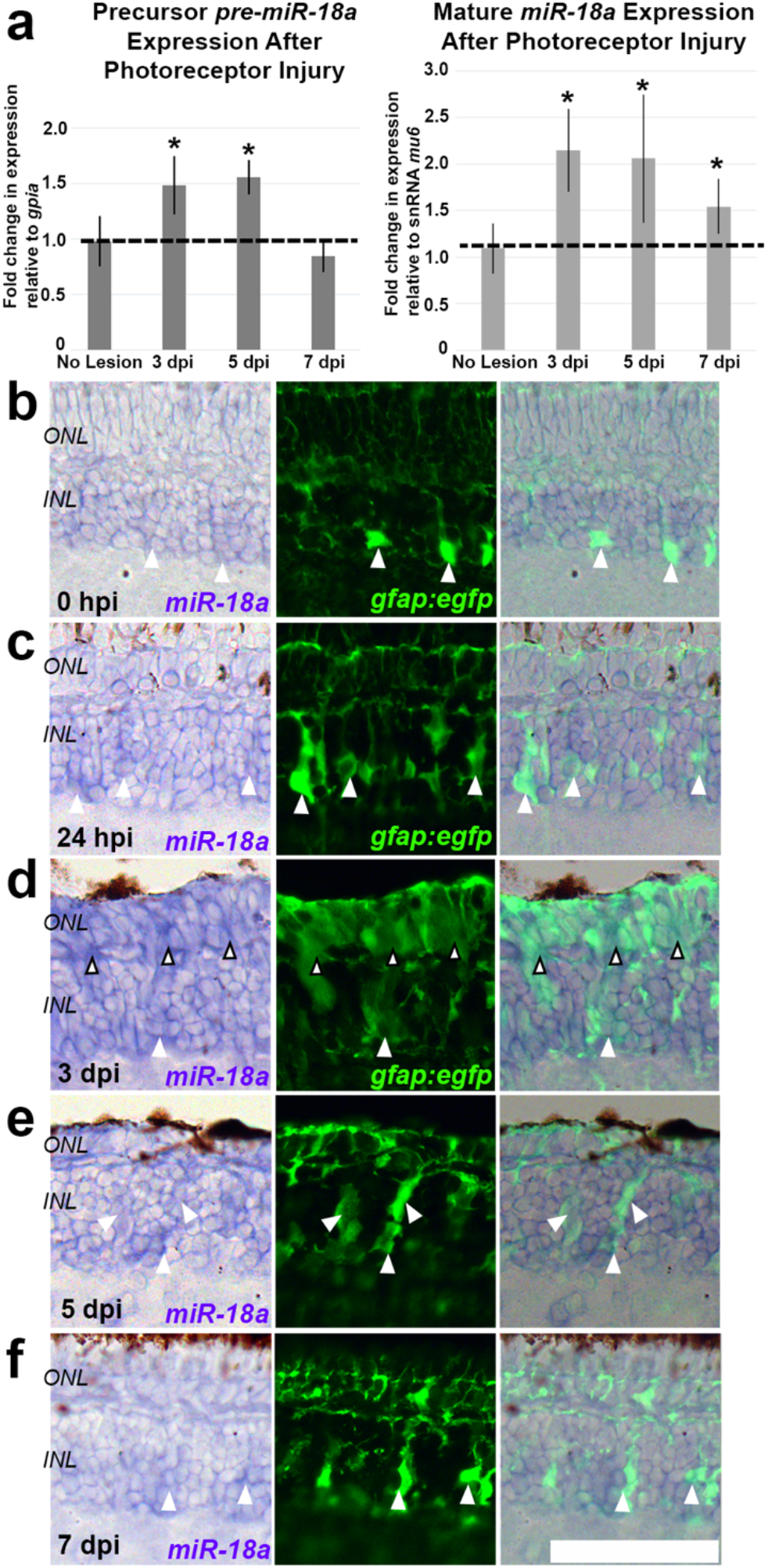
Expression of *miR-18a* in the retina following photoreceptor injury. (a) RT-qPCR showing retinal expression of the precursor for *miR-18a* (*pre-mir-18a*) with higher expression than controls (no lesion) at 3 days (1.48 ± 0.26 SD fold higher, p=0.021, n=4) and 5 days post-injury (dpi) (1.56 ± 0.16 SD fold higher, p=0.007, n=4). (b) Taqman RT-qPCR showing retinal expression of *miR-18a* in control (no lesion) retinas compared with 3 dpi (2.15 ± 0.44 SD fold higher, p=0.003, n=5), 5 dpi (2.06 ± 0.69 SD fold higher, p=0.0.024, n=5) and 7 dpi (1.54 ± 0.30 SD fold higher, p=0.045, n=5). Expression (fold differences) are calculated using the DDCT method, relative to housekeeping gene *gpia* for (a) and the small nuclear protein RNA U6 (*mu6*) for (b). Error bars represent standard deviation and asterisks indicate significant differences between each injury time point and controls (Student’s t-test, p<0.05). (b-f) *In situ* hybridizations for *miR-18a* (purple) in retinal cross sections at different post-injury time points in *Tg(gfap:egfp)^mi2002^* fish, in which Müller glia and MG-derived progenitors express *egfp*, visualized with immunolabeling for EGFP protein (green). White arrowheads show examples of EGFP-labeled Müller glia that express *miR-18a* and black outlined arrowheads show examples of MG-derived photoreceptor progenitors in the ONL that express *miR-18a*. Abbreviations: ONL—outer nuclear layer, INL—inner nuclear layer; *scale bar:* 50 mm

**Fig. 2.**
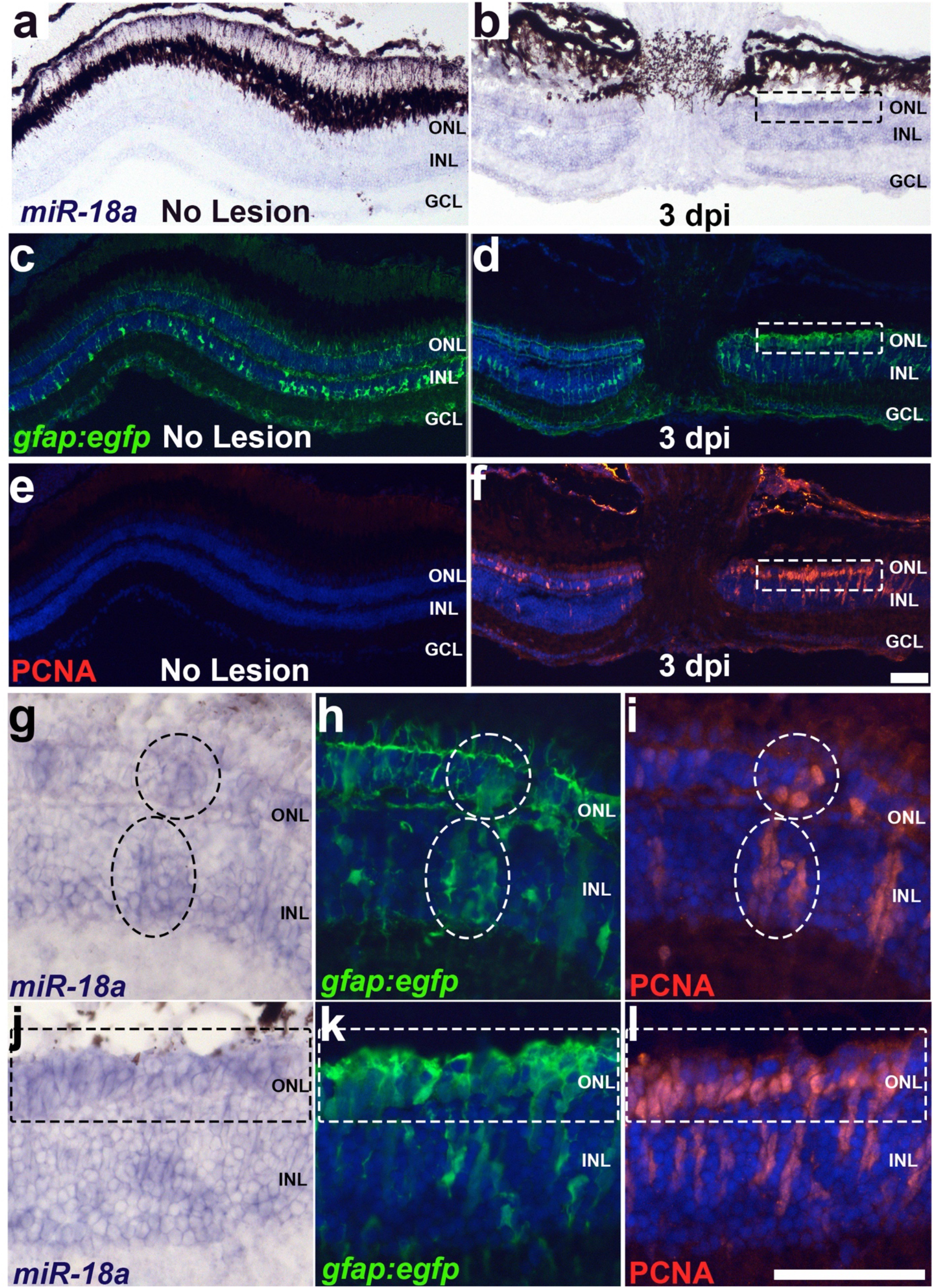
*In situ* hybridizations for *miR-18a* (purple) (a, b) in retinal cross sections at 3 dpi compared with uninjured retinas in *Tg(gfap:egfp)^mi2002^* fish, in which Müller glia and MG-derived progenitors express *egfp*, visualized with immunolabeling for GFP protein (green) (c, d, h, k). Proliferating cells are immunolabeled with anti-proliferating cell nuclear antigen (PCNA) (red) (e, f, i, l). Clusters of cells surrounded by dotted boxes or elipses indicate co-labeled cells. Abbreviations: ONL—outer nuclear layer, INL—inner nuclear layer, GCL—ganglion cell layer; *scale bar:* 50 μm

### Loss of *miR-18a* alters the timing, but not the extent, of photoreceptor regeneration

In the embryonic zebrafish retina, the absence of *miR-18a* results in accelerated photoreceptor differentiation but, by 6 days post-fertilization, does not affect the number of photoreceptors produced [30]. Previous work did not compare photoreceptor numbers between WT and *miR-18a^mi5012^* fish in the adult, uninjured retina. To determine this, immunolabeling was used to label mature cones (Zpr-1) and rods (Zpr-3) in uninjured WT and *miR-18a^mi5012^* retinas, and this showed that there are no differences in photoreceptor numbers in the uninjured retina (Figure 3a, b, i; Figure 4a, b, i). Further, *in situ* hybridization, used to label mature cones (*arr3a*) and rods (*rho*), also showed no difference (Figure S1a, b, i, j).

**Fig. 3.**
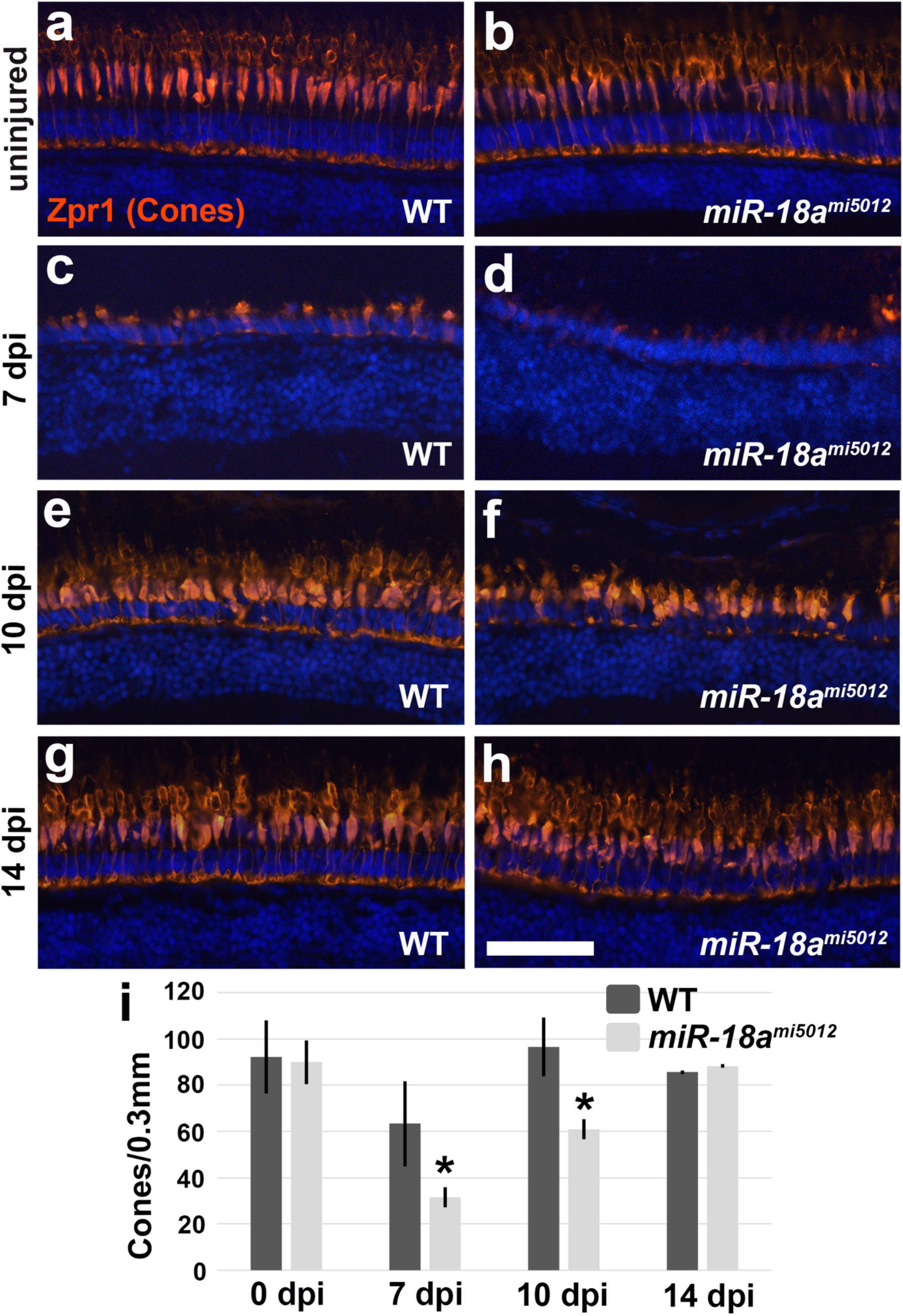
Comparison and quantification of mature cone photoreceptors in WT and *miR-18a^mi5012^* retinas at uninjured (a, b), 7 dpi (c, d), 10 dpi (e, f) and 14 dpi (g, h), and quantified in (i), using immunolabeling for cones (Zpr-1/Arr3; red immunolabeling). Photoreceptor counts in retinal cross sections in the center of the lesioned area (cells per 0.3 mm of linear retina). Counts are as follows: uninjured (WT 92.1 ± 15.8 SD, *miR-18a^mi5012^* 89.9 ± 9.4 SD cells/0.3 mm, p=0.976, n=3), 7 dpi (WT 63.3 ± 18.4 SD, *miR-18a^mi5012^* 31.4 ± 4.3 SD cells/0.3 mm, p=0.017, n=3), 10 dpi (WT 96.4 ± 12.6 SD, *miR-18a^mi5012^* 60.9 ± 4.2 SD cells/0.3 mm, p=0.005, n=3), and 14 dpi (WT 85.6 ± 0.77 SD, *miR-18a^mi5012^* 88.3 ± 0.84 SD cells/0.3 mm, p=0.137, n=3). Error bars represent standard deviation and asterisks indicate significant differences (Student’s t-test, p<0.05). Abbreviations: ONL—outer nuclear layer, INL—inner nuclear layer; *scale bar:* 50 mm

**Fig. 4.**
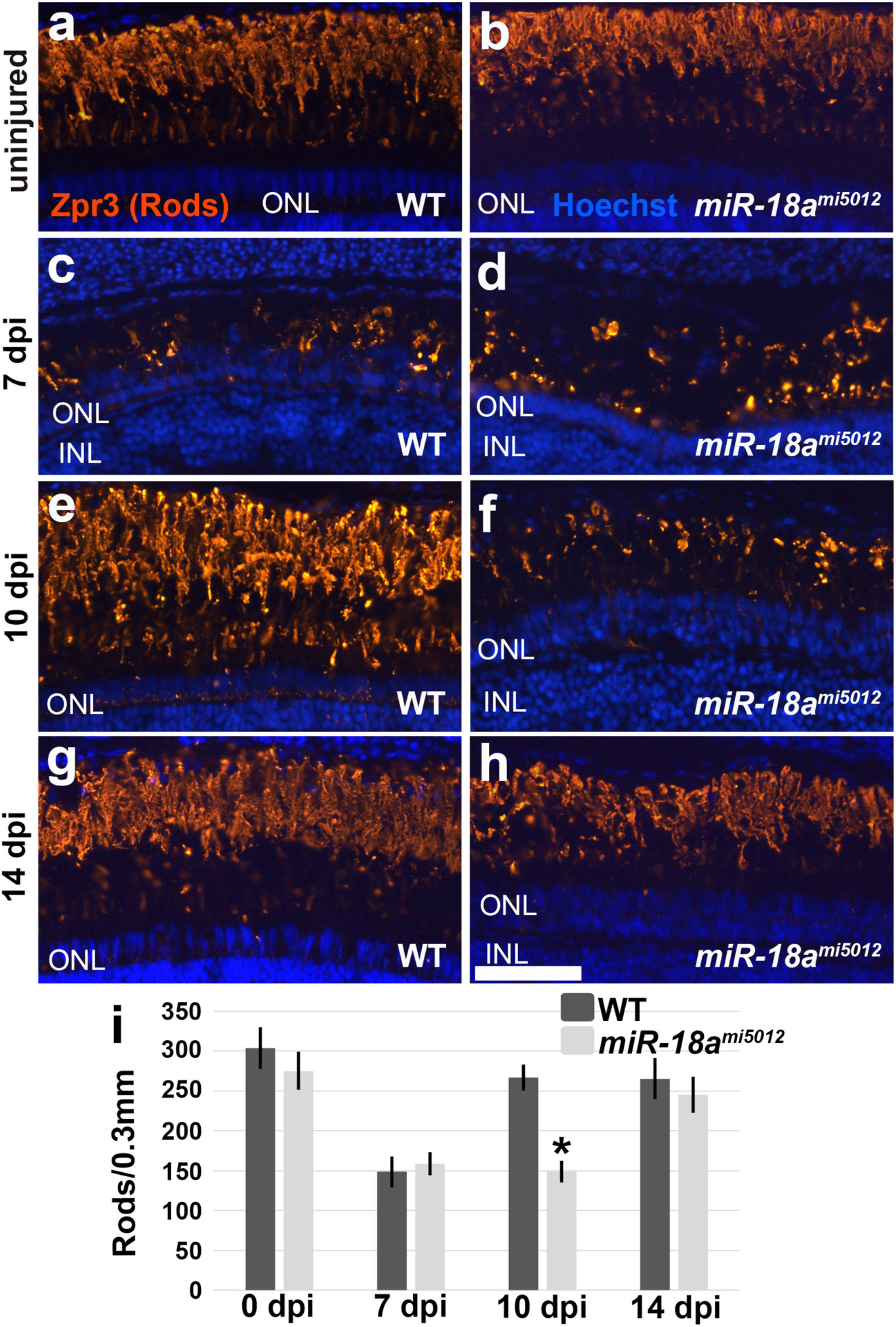
Comparison and quantification of mature rod photoreceptors in WT and *miR-18a^mi5012^* retinas at uninjured (a, b), 7 dpi (c, d), 10 dpi (e, f) and 14 dpi (g, h), and quantified in (i), using immunolabeling for rods (Zpr-3/Rho; red immunolabeling).. Photoreceptor counts in retinal cross sections in the center of the lesioned area (cells per 0.3 mm of linear retina). Counts are as follows: uninjured (WT 304.0 ± 26.6 SD, *miR-18a^mi5012^* 275.6 ± 24.1 SD cells/0.3 mm, p=0.242, n=3), 7 dpi (WT 148.6 ± 19.2 SD, *miR-18a^mi5012^* 158.7 ± 14.1 SD cells/0.3 mm, p=0.499, n=3), 10 dpi (WT 266.8 ± 16.2 SD, *miR-18a^mi5012^* 148.8 ± 13.2 SD cells/0.3 mm, p<0.001, n=3), and 14 dpi (WT 265.6 ± 26.0 SD, *miR-18a^mi5012^* 245.4 ± 22.5 SD cells/0.3 mm, p=0.369, n=3). Error bars represent standard deviation and asterisks indicate significant differences (Student’s t-test, p<0.05). Abbreviations: ONL—outer nuclear layer, INL—inner nuclear layer; *scale bar:* 50 mm

To determine if the absence of *miR-18a* affects the timing and/or extent of photoreceptor regeneration in adults, immunolabeling and *in situ* hybridization were used to label mature cones and rods in injured retinas. First, mature photoreceptors were quantified in WT retinas at 3 dpi, to establish a baseline for comparison and determine the numbers of photoreceptors destroyed by the photolytic lesioning method. This determined that at 3 dpi, in the central retina just dorsal to the optic nerve, 96.4% of cones (Figure S2a, c, e) and 61.6% of rods (Figure S2b, d, f) had been destroyed. Next, quantitative comparisons were made between WT and *miR-18a^mi5012^* retinas at 7 dpi, when large numbers of newly differentiated photoreceptors can first be detected, 10 dpi, when most new photoreceptors have been normally regenerated, and 14 dpi, when photoreceptor regeneration is largely complete. At 7 dpi, *miR-18a^mi5012^* retinas have fewer mature cones than WT (Figure 3c, d, i), but the number of mature rods does not differ (Figure 4c, d, i). At 10 dpi, *miR-18a* retinas have fewer mature cones and rods than WT (Figure 3e, f, i; Figure 4e, f, i) but at 14 dpi, the number of mature photoreceptors does not differ (Figure 3g-i; Figure 4g-i). To confirm the differences in photoreceptor numbers detected with immunohistochemistry, *in situ* hybridization was used to label cones (*arr3a*) and rods (*rho*) at the same time points (Figure S1). These data confirmed that, compared with WT, *miR-18a* retinas have fewer mature cones at 7 dpi and fewer mature cones and rods at 10 dpi. Together, these data show that in *miR-18a^mi5012^* retinas compared with WT, the same overall numbers of photoreceptors are present prior to injury and are regenerated by 14 dpi, but maturation of cone and rod photoreceptors is delayed.

### During photoreceptor regeneration, the absence of *miR-18a* results in hyperproliferation among Müller glia-derived progenitors

The delay in photoreceptor maturation in the injured *miR-18a^mi5012^* retina indicates that photoreceptor regeneration is delayed, and this could be linked to the timing of proliferation among photoreceptor progenitors. To determine if *miR-18a* regulates cell proliferation among MG-derived progenitors, BrdU labeling and immunostaining were used to quantitatively compare the number of proliferating cells in WT and *miR-18a^mi5012^* retinas. In the injured WT retina, Müller glia divide between 1 and 2 dpi to produce neural progenitors, MG-derived progenitors then divide multiple times with proliferation peaking around 3 dpi, many MG-derived progenitors normally stop dividing between 4 and 5 dpi, and the first regenerated photoreceptors can be detected between 5 and 6 dpi [see 42]. By 7 and 10 dpi, proliferation is markedly reduced, and very few progenitors normally continue to proliferate. Compared with WT retinas, the total number of BrdU-labeled cells [i.e. cells that were actively dividing during the 24-hour period of BrdU exposure] in *miR-18a^mi5012^* retinas did not differ at 3 dpi (Figure 5a, b), and there were no differences in numbers of BrdU-labeled cells in either the INL (WT 50.0 ± 7.2 SD, *miR-18a^mi5012^* 40 ± 10.8 SD cells/0.3 mm, p=0.25, n=3) or ONL (WT 17.7 ± 3.8 SD, *miR-18a^mi5012^* 14.2 ± 3.2 SD cells/0.3 mm, p=0.29, n=3). This indicates that the initial proliferative response is unaltered in *miR-18a^mi5012^* retinas. However, at 7 dpi, there were significantly more BrdU-labeled cells in the *miR-18a^mi5012^* retinas than in WT (Figure 5c, d), and this difference was most pronounced in the ONL (WT 21.8 ± 5.9 SD, *miR-18a^mi5012^* 51.4 ± 6.7 SD cells/0.3 mm, p=0.0003, n=3). At 10 dpi, there were also significantly more BrdU-labeled cells in the *miR-18a^mi5012^* retinas (Figure 5e, f), and significantly more cells in both the INL (WT 1.7 ± 0.8 SD, *miR-18a^mi5012^* 14.3 ± 2.5 SD cells/0.3 mm, p=0.001, n=3) and ONL (WT 11.0 ± 1.8 SD, *miR-18a^mi5012^* 30.2 ± 4.9 SD cells/0.3 mm, p=0.003, n=3).

**Fig. 5.**
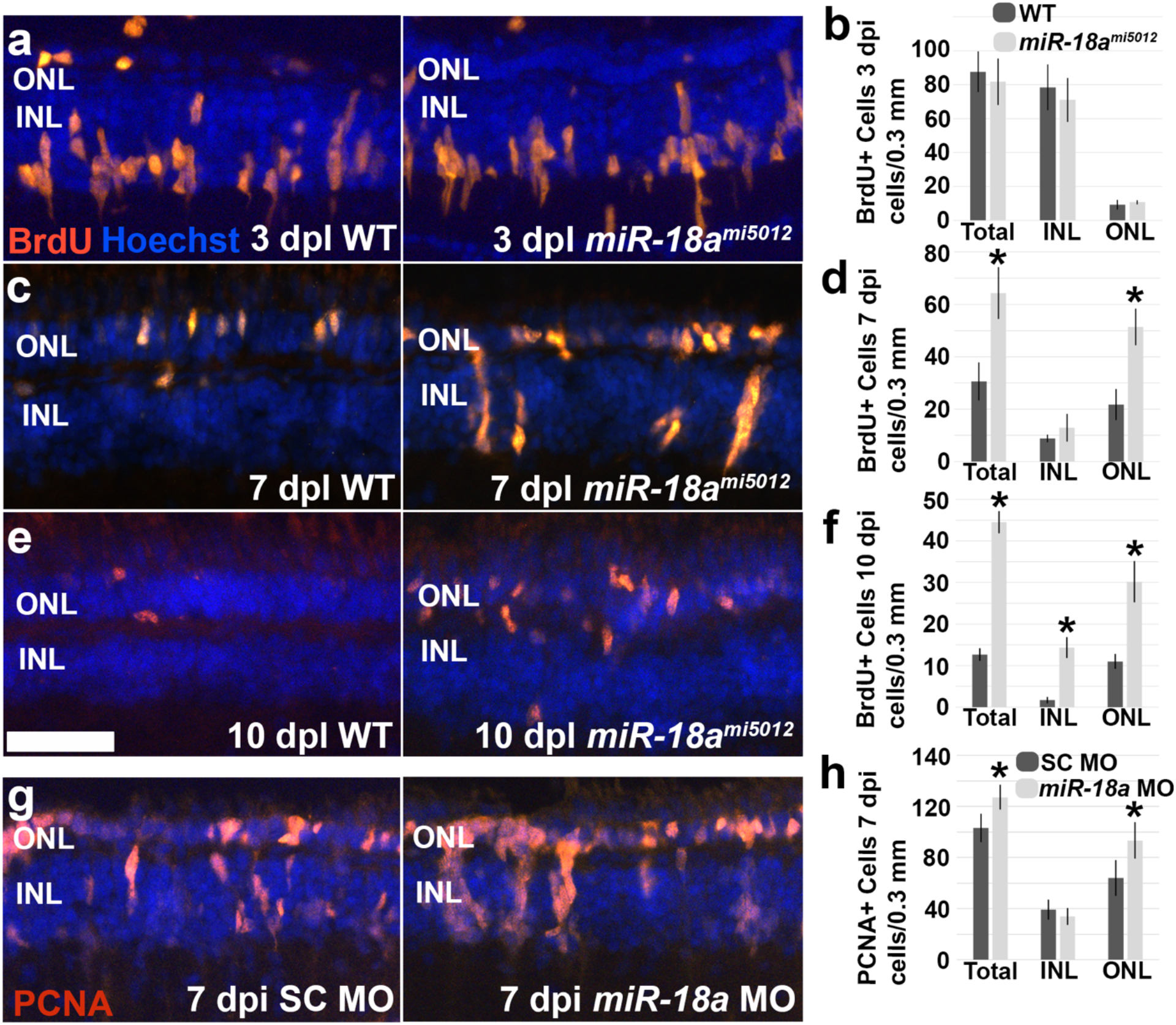
Comparison of the numbers of proliferating cells in WT and *miR-18a^mi5012^* retinas at 3, 7 and 10 days post-photoreceptor injury (dpi), and in control morpholino (MO) and *miR-18a* MO injected retinas at 7 dpi. (a) BrdU immunolabeling in 3 dpi retinas in which fish were immersed in 5 mM BrdU from 2 to 3 dpi; BrdU-labeled cells (S phase of the cell cycle) are shown in orange-red and Hoechst labeled cell nuclei are blue (b) BrdU-labeled cell counts in these retinas in cells per 0.3 mm of linear retina (WT 87.6 ± 8.1 SD, *miR-18a^mi5012^* 81.7 ± 7.2 SD cells/0.3 mm, p=0.60, n=3). (c) BrdU immunolabeling in 7 dpi retinas in which fish were immersed in 5 mM BrdU from 6 to 7 dpi; and (d) BrdU-labeled cell counts in these retinas in cells per 0.3 mm of linear retina (WT 30.6 ± 7.2 SD, *miR-18a^mi5012^* 64.3 ± 9.8 SD cells/0.3 mm, p=0.001, n=3). (e) BrdU immunolabeling in 10 dpi retinas in which fish were immersed in 5 mM BrdU from 9 to 10 dpi; and (f) BrdU-labeled cell counts in these retinas in cells per 0.3 mm of linear retina (WT 12.7 ± 1.5 SD, *miR-18a^mi5012^* 44.5 ± 2.6 SD cells/0.3 mm, p<0.0001, n=3). (g) PCNA immunolabeling in 7 dpi retinas from eyes that were injected/electroporated with standard control morpholinos (SC MO) or *miR-18a* MO at 2 dpi; and (h) PCNA-immunolabeled cell counts in these retinas in cells per 0.3 mm of linear retina (total PCNA+ cells: SC MO 103.0 ± 11.1 SD, *miR-18a* MO 127.1 ± 9.6 SD cells/0.3 mm, p<0.004, n=5; ONL PCNA+ cells: SC MO 63.9 ± 14.1 SD, *miR-18a* MO 93.4 ± 14.2 SD cells/0.3 mm, p<0.008, n=5). Error bars represent standard deviation and asterisks indicate significant differences (Student’s t-test, p<0.05); Abbreviations: ONL—outer nuclear layer, INL—inner nuclear layer; *scale bar:* 50 mm

To confirm the excess proliferation observed for *miR-18a^mi5012^* retinas, a second, independent approach was used to deplete *miR-18a* during photoreceptor regeneration. A *miR-18a* morpholino oligonucleotide, previously confirmed to effectively knock down *miR-18a* [38, 30], was used to knock down *miR-18a* in WT retinas during photoreceptor regeneration. Because *miR-18a* expression peaks at 3 dpi, for the present experiment, morpholinos were injected and electroporated at 2 dpi and eyes collected at 7 dpi. Compared with a standard control morpholino (SC MO), at 7 dpi, *miR-18a* knockdown resulted in more PCNA-immunolabeled progenitor cells [i.e proliferating cells that are in G1 to S-phase of the cell cycle] in the retina and, as in *miR-18a^mi5012^* retinas, there were more dividing progenitors specifically in the ONL (Figure 5g, h). The MO injection and electroporation technique is invasive and could possibly increase the overall proliferative response; PCNA might also label more cells than the 24-hour BrdU exposure; together, these differences likely explain the higher overall numbers of PCNA-labeled cells observed in MO injected compared with BrdU-labeled cells in 7 dpi uninjected retinas (compare with Figure 5c, d). Together, along with *miR-18a^mi5012^* data, these results show that following photoreceptor injury, in the absence of *miR-18a*, the timing of the initial proliferative response is unchanged, but that progenitors continue to proliferate longer than in WT retinas.

### Following photoreceptor injury in miR-18a^mi5012^ retinas, excess progenitors migrate to all retinal layers

In *miR-18a^mi5012^* retinas, MG-derived progenitors proliferate for a longer period of time, suggesting that excess MG-derived progenitors are produced. However, since extra photoreceptors are not generated, the excess progenitors might either die or migrate to other retinal layers, possibly differentiating into other cell types. To first determine the proportion of progenitors that differentiate into photoreceptors, BrdU tracing was performed by exposing fish to BrdU from 3 to 5 dpi during the peak of cell proliferation and then, at 7 and 10 dpi, performing BrdU immunolabeling combined with *in situ* hybridization for cones and rods. The results showed that at both 7 and 10 dpi compared with WT, *miR-18a^mi5012^* retinas had more BrdU-positive cells (Figure 6a-e, 7a-e), showing that, between 3 and 5 dpi, mutant retinas generate more dividing progenitors. At 7 dpi compared with WT, a lower percentage of total BrdU-positive cells in *miR-18a^mi5012^* retinas had migrated to the ONL and, of the progenitors that had migrated to the ONL, a lower percentage had differentiated into cone photoreceptors (Figure 6a, b, f). At 10 dpi the percentage of total BrdU-positive cells in the ONL was still lower in *miR-18a^mi5012^* retinas than in WT, and so were the percentages of BrdU-positive cells in the ONL that had differentiated into both cones and rods (Figure 7a-d, f). Additionally in *miR-18a^mi5012^* retinas, greater percentages of the BrdU-positive cells were present in the INL and the ganglion cell layer (GCL) (Figure 7a-d, f). Together, these data show that in *miR-18a^mi5012^* retinas, progenitors remain in the inner retina longer than in WT retinas and, of the progenitors that migrate to the ONL by 7 or 10 dpi, fewer differentiate into photoreceptors.

**Fig. 6.**
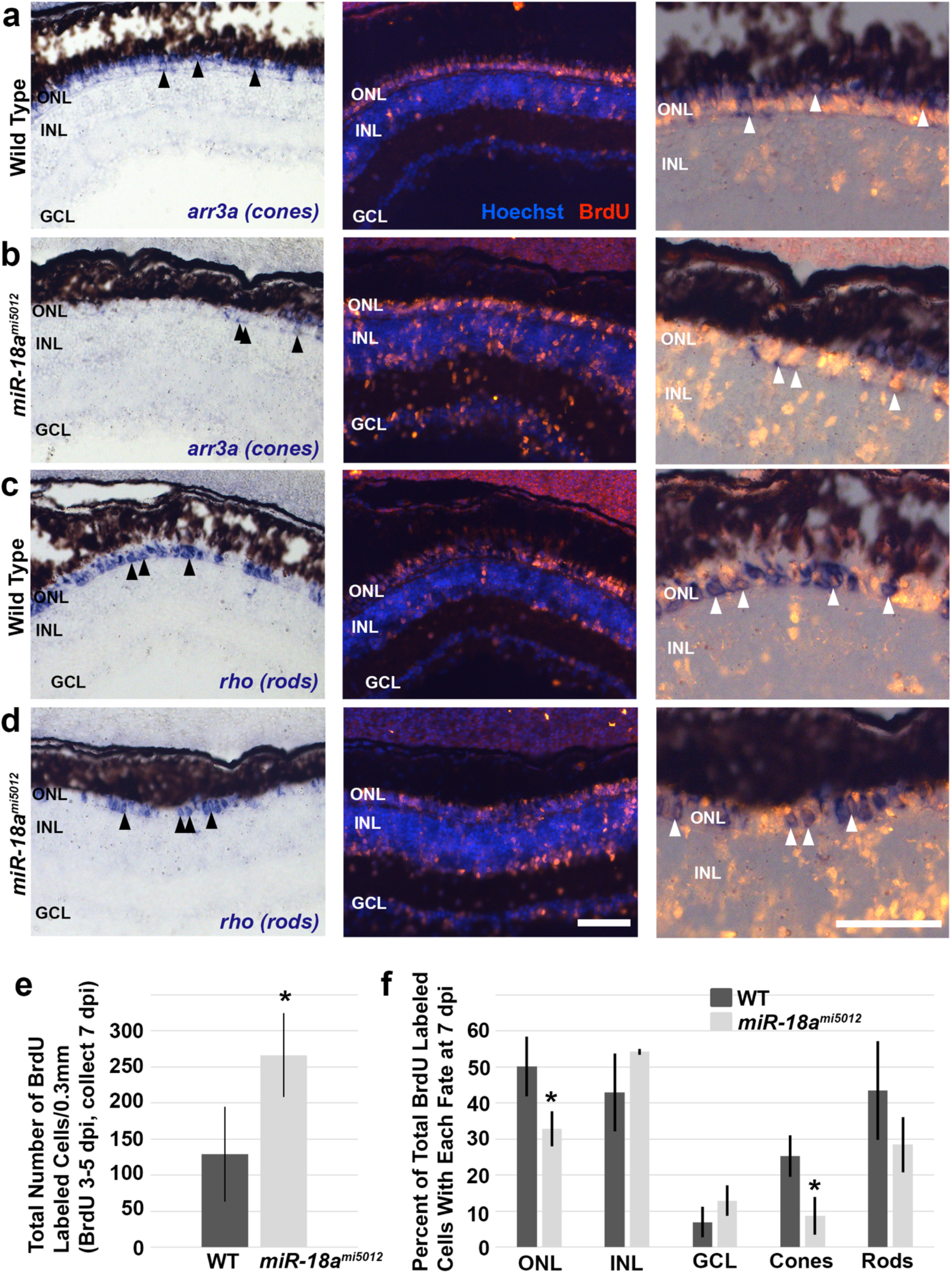
In situ hybridization for mature cone and rod photoreceptors at 7 dpi combined with BrdU immunolabeling in retinas of WT and *miR-18a^mi5012^* fish exposed to BrdU from 3 to 5 dpi. (a, b) *In situ* hybridization for cones (*arr3a*, left panels) where black arrowheads show examples of labeled mature cones, BrdU immunolabeling (center panels in red), and combined *in situ* hybridization and BrdU immunolabeling (right panels) where white arrowheads show examples of co-labeled cells that are newly regenerated cones. (c, d) *In situ* hybridization for rods (*rho*, left panels) where black arrowheads show examples of labeled mature rods, BrdU immunolabeling (center panels in red), and combined *in situ* hybridization and BrdU immunolabeling (right panels) where white arrowheads show examples of co-labeled cells that are newly regenerated rods. (e) Quantification of total numbers of BrdU labeled cells (WT 128.7 ± 65.9 SD, *miR-18a^mi5012^* 266.6 ± 58.5 SD cells/0.3 mm, p=0.027, n=3), and (f) quantification of the percentage of total BrdU labeled cells that are in the outer nuclear layer (ONL) (WT 50.1% ± 8.3 SD, *miR-18a^mi5012^* 32.9% ± 4.9 SD, p=0.018, n=3), inner nuclear layer (INL) (WT 42.9% ± 10.8 SD, *miR-18a^mi5012^* 52.3% ± 0.8 SD, p=0.55, n=3), ganglion cell layer (GCL) (WT 6.8% ± 4.2 SD, *miR-18a^mi5012^* 12.8% ± 4.2 SD, p=0.16, n=3), and that have differentiated into cones (WT 25.2% ± 5.7 SD, *miR-18a^mi5012^* 8.6% ± 5.3 SD, p=0.011, n=3) or rods (WT 43.5% ± 13.7 SD, *miR-18a^mi5012^* 28.4% ± 7.7 SD, p=0.17, n=3) (co-label with the *in situ* hybridization marker). Cells were counted in retinal cross sections across 0.3 mm of linear retina. Error bars represent standard deviation and asterisks indicate significant differences (Student’s t-test, p<0.05); Abbreviations: ONL—outer nuclear layer, INL—inner nuclear layer, GCL—ganglion cell layer; *scale bars:* 50 μm

**Fig. 7.**
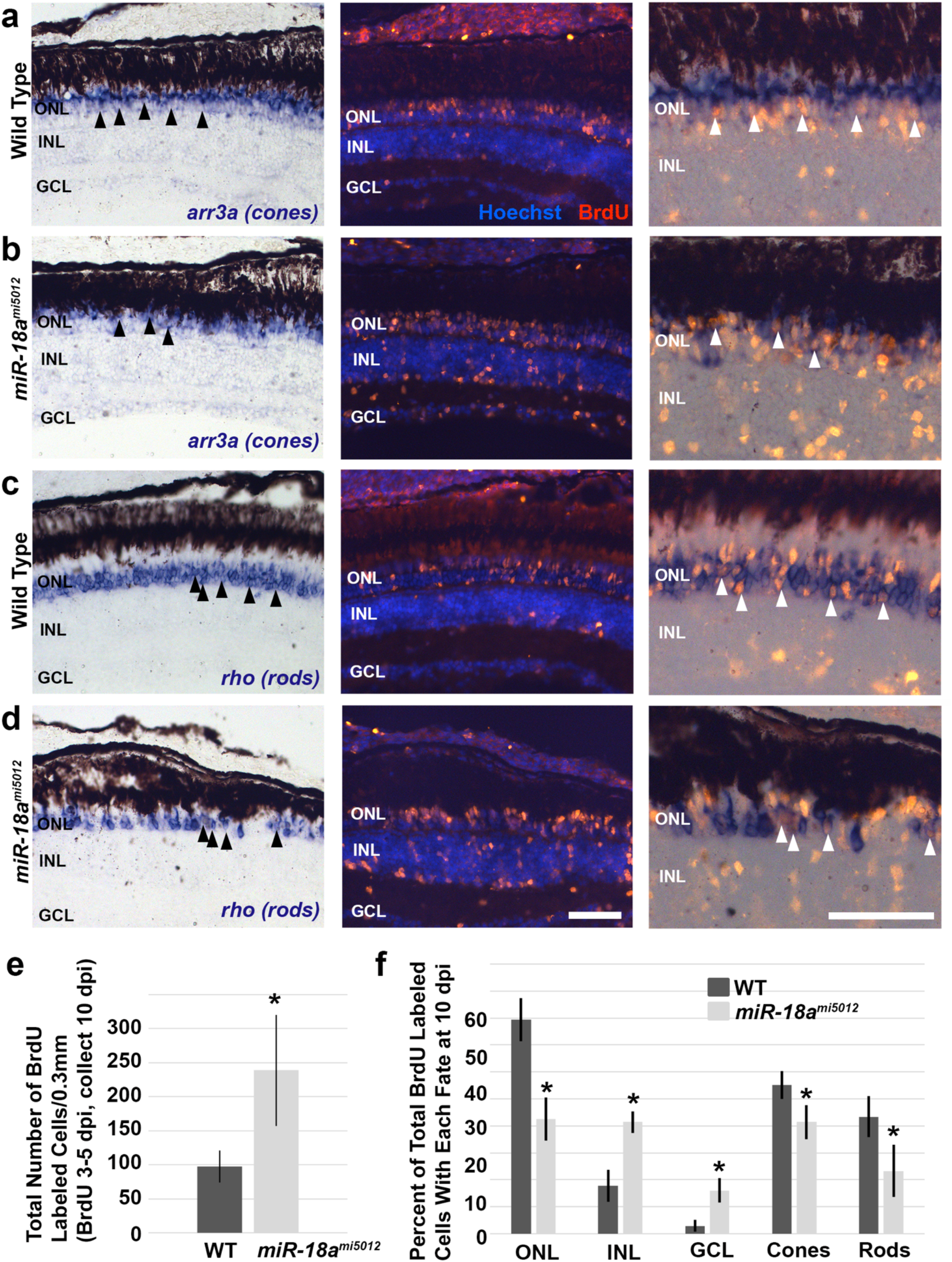
In situ hybridization for mature cone and rod photoreceptors at 10 dpi combined with BrdU immunolabeling in retinas of WT and *miR-18a^mi5012^* fish exposed to BrdU from 3 to 5 dpi. (a, b) *In situ* hybridization for cones (*arr3a*-left panels) where black arrowheads show examples of labeled mature cones, BrdU immunolabeling (center panels in red), and combined *in situ* hybridization and BrdU immunolabeling (right panels) where white arrowheads show examples of co-labeled cells that are newly regenerated cones. (c, d) *In situ* hybridization for rods (*rho*-left panels) where black arrowheads show examples of labeled mature rods, BrdU immunolabeling (center panels in red), and combined *in situ* hybridization and BrdU immunolabeling (right panels) where white arrowheads show examples of co-labeled cells that are newly regenerated rods. (e) Quantification of total numbers of BrdU labeled cells (WT 97.8 ± 23.4 SD, *miR-18a^mi5012^* 238.7 ± 81.5 SD cells/0.3 mm, p=0.023, n=3), and (f) quantification of the percentage of total BrdU labeled cells that are in the outer nuclear layer (ONL) (WT 79.3% ± 8.0 SD, *miR-18a^mi5012^* 42.6% ± 8.0 SD, p=0.002, n=3), inner nuclear layer (INL) (WT 17.8% ± 5.9 SD, *miR-18a^mi5012^* 41.4% ± 3.9 SD, p=0.002, n=3), ganglion cell layer (GCL) (WT 2.9% ± 2.1 SD, *miR-18a^mi5012^* 16.1% ± 4.5 SD, p=0.005, n=3), and that have differentiated into cones (WT 55.2% ± 5.1 SD, *miR-18a^mi5012^* 41.4% ± 6.3 SD, p=0.021, n=3) or rods (WT 43.4% ± 7.6 SD, *miR-18a^mi5012^* 23.3% ± 9.7 SD, p=0.024, n=3) (co-label with the *in situ* hybridization marker). Cells were counted in retinal cross sections across 0.3 mm of linear retina. Error bars represent standard deviation and asterisks indicate significant differences (Student’s t-test, p<0.05); Abbreviations: ONL—outer nuclear layer, INL—inner nuclear layer, GCL—ganglion cell layer; *scale bars:* 50 μm

To determine if the excess progenitors in *miR-18a^mi5012^* retinas survive, TUNEL was used to label apoptotic cells at 10 dpi, when the largest differences in numbers of photoreceptors and MG-derived progenitors were identified between WT and *miR-18a^mi5012^* retinas. This experiment showed that, at 10 dpi, there were no TUNEL-positive cells in either WT or *miR-18a^mi5012^* retinas (Figure 8a), although some TUNEL-positive cells were observed in extraocular tissues in both WT and *miR-18a^mi5012^* (arrowheads, Figure 8a), thereby, demonstrating the sensitivity of the assay.

**Fig. 8.**
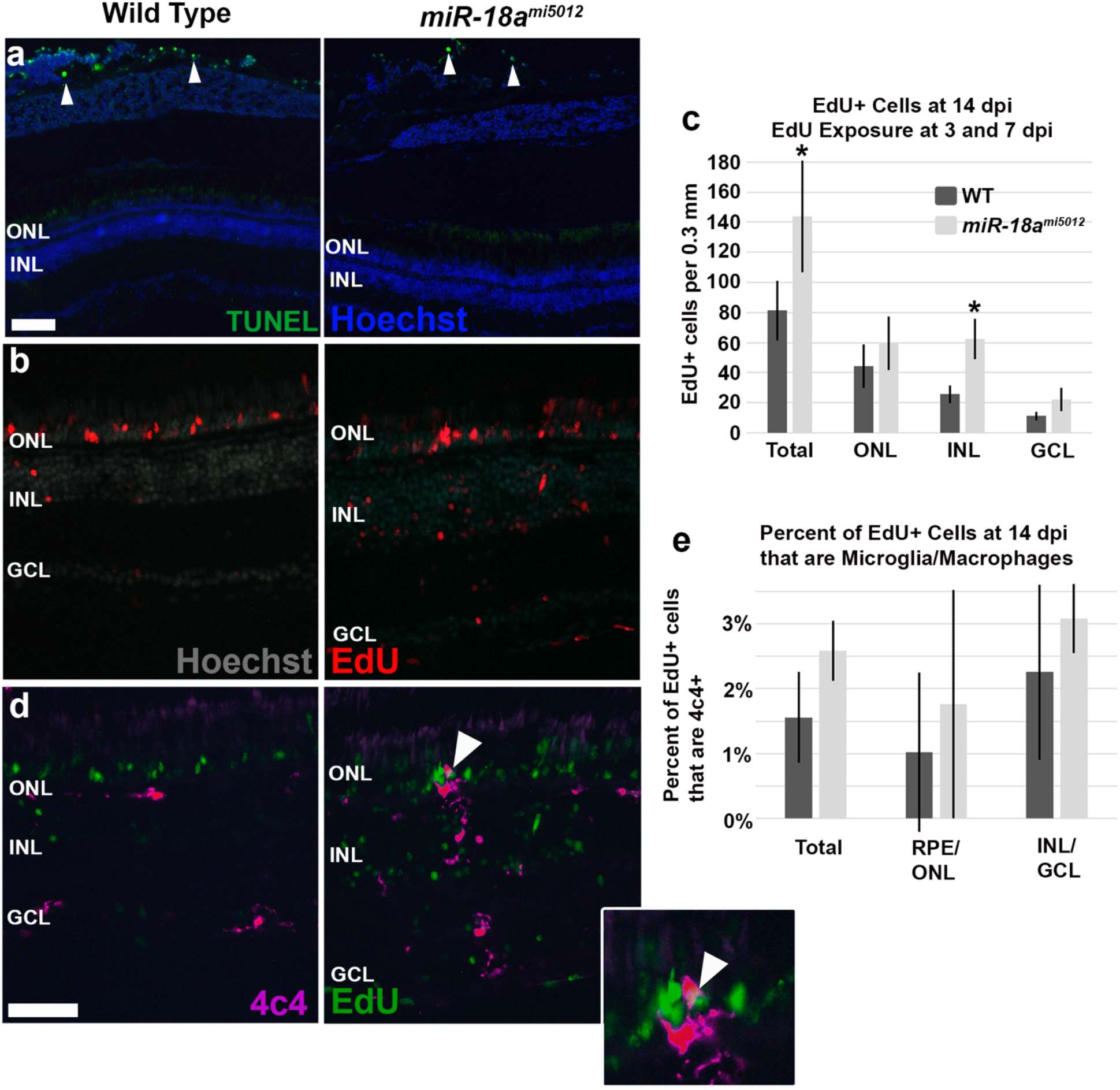
The destinations of progenitor cells following photoreceptor injury in 14 dpi WT and *miR-18a^mi5012^* retinas from fish that were injected with EdU at 3 and 7 dpi. (a) TUNEL labeling at 10 dpi showing no apoptotic cells in WT or *miR-18a^mi5012^* retinas, but extraocular tissues have some TUNEL-positive cells (arrowheads). (b) EdU labeling of WT and *miR-18a^mi5012^* retinal cells at 14 dpi, from fish that were exposed to EdU (IP injection) at 3 and 7 dpi. (c) EdU-positive cells (red) are quantified in total (WT 81.2 ± 19.7 SD, *miR-18a^mi5012^* 144.0 ± 37.1 SD cells/0.3 mm, p=0.047, n=4), in the outer nuclear layer (ONL—WT 44.4 ± 14.4 SD, *miR-18a^mi5012^* 59.4 ± 17.8 SD cells/0.3 mm, p=0.290, n=4), inner nuclear layer (INL—WT 25.7 ± 6.0 SD, *miR-18a^mi5012^* 62.3 ± 13.3 SD cells/0.3 mm, p=0.007, n=4) and in/near the ganglion cell layer (GCL—WT 11.1 ± 3.0 SD, *miR-18a^mi5012^* 22.3 ± 7.7 SD cells/0.3 mm, p=0.066, n=4). (d) Edu labeling (green) combined with 4c4 immunolabeling of microglia/macrophages (magenta) showing rare EdU/4c4 co-labeled cells (arrowhead—enlarged in the adjacent box). (e) Percent of EdU-labeled cells at 14 dpi (injections at 3 and 7 dpi) that are co-labeled with 4c4 (total—WT 1.6% ± 0.7% SD, *miR-18a^mi5012^* 2.6% ± 0.5% SD cells/0.3 mm, p=0.066, n=4; RPE/ONL—WT 1.0% ± 1.2% SD, *miR-18a^mi5012^* 1.8% ± 1.8% SD cells/0.3 mm, p=0.484, n=4; INL/GCL—WT 2.3% ± 1.4% SD, *miR-18a^mi5012^* 3.1% ± 0.5% SD cells/0.3 mm, p=0.305, n=4). Cells were counted in retinal cross sections across 0.3 mm of linear retina. Error bars represent standard deviation and asterisks indicate significant differences (Student’s t-test, p<0.05); *scale bar:* 50 μm. Abbreviations: ONL—outer nuclear layer, INL—inner nuclear layer; *scale bar:* 50 μm

The absence of TUNEL-positive cells in the retinas of WT and mutants indicates that excess progenitors in *miR-18a^mi5012^* retinas are not eliminated by cell death. To determine if excess progenitors in *miR-18a^mi5012^* retinas remain in the inner retinal layers, fish were injected with 1 mg/L EdU at 3 and 7 dpi, corresponding to times when proliferation peaks (3 dpi) and when significanty more proliferating cells are present in *miR-18a^mi5012^* retinas. Fish were then sacrificed at 14 dpi when photoreceptor regeneration is complete and total photoreceptor numbers no longer differ between *miR-18a^mi5012^* and WT. EdU visualization at 14 dpi showed that, compared with WT, *miR-18a^mi5012^* retinas have more EdU-labeled cells in the retina, and particularly in the inner nuclear layer (INL) (Figure 8b, c), indicating that many of the excess progenitors produced at 3-7 dpi remain in the INL and, by 14 dpi, do not adopt a photoreceptor fate.

One possible fate of the excess proliferating cells produced in the injured *miR-18a^mi5012^* retina is that some might differentate into microglia/macrophage cells that play a critical role in the post-injury and inflammatory response. To determine this, retinal sections from fish injected with EdU a 3 and 7 dpi were immunolabeled with an antibody specific for microglia/macrophage cells (4c4), and this was followed by visualization of EdU. The results showed that in both WT and *miR-18a^mi5012^* retinas at 14 dpi, only a very small (and not significantly different) percentage of the EdU-positive cells were co-labeled with the 4c4 antibody (Figure 8d, e). This indicates that most of the excess progenitors produced in injured *miR-18a^mi5012^* retinas do not differentiate into microglia/macrophage cells.

### During photoreceptor regeneration, lack of *miR-18a* results in prolonged inflammation

The effect of *miR-18a* on the cell cycle during photoreceptor regeneration is strikingly different from embryonic development, in which *miR-18a* regulates photoreceptor differentiation but not cell proliferation [30]. This indicates that in the injured retina, *miR-18a* regulates pathways that are specific to the post-injury response. Silva *et al.* [21] showed that in injured *mmp9* mutant retinas, there was increased inflammation resulting in excess proliferation among MG-derived progenitors. This finding helped lead to the rationale that increased or prolonged inflammation might cause the excess proliferation observed in *miR-18a^mi5012^* retinas. Indeed, the miRNA target database TargetscanFish (http://www.targetscan.org/fish_62) predicts that *miR-18a* interacts with mRNA for many molecules involved in inflammatory pathways. To determine if *miR-18a* regulates inflammation, RT-qPCR was used to compare the mRNA expression levels of key inflammatory molecules at 1, 3, 5 and 7 dpi in WT and *miR-18a^mi5012^* retinas. These data show that at 1 dpi, around the time that Müller glia normally begin to divide, expression of *tnfα* is higher in *miR-18a^mi5012^* retinas compared with WT (WT 2.31 ± 0.99 SD, *miR-18a^mi5012^* 4.51 ± 1.14 SD, p=0.033, n=3) (Figure 9a). Then at 3 dpi, around the normal peak of MG-derived progenitor proliferation, *il1β* expression is higher in *miR-18a^mi5012^* retinas compared with WT (WT 0.97 ± 0.06 SD, *miR-18a^mi5012^* 1.61 ± 0.21 SD, p=0.007, n=3) (Figure 9b). At 5 dpi, when many photoreceptor progenitors normally stop proliferating and begin to differentiate, expression of *il6*, *il1β*, and the cytokine regulator Nuclear Factor Kappa B 1 (*nfkb1*) are higher in *miR-18a^mi5012^* retinas compared with WT (*il6* WT 0.85 ± 0.25 SD, *miR-18a^mi5012^* 1.32 ± 0.26 SD, p=0.042, n=3; *il1b* WT 0.97 ± 0.08 SD, *miR-18a^mi5012^* 1.71 ± 0.47 SD, p=0.028, n=3; *nfkb1* WT 1.19 ± 0.17 SD, *miR-18a^mi5012^* 1.68 ± 0.23 SD, p=0.021, n=3) (Figure 9c). Finally, at 7 dpi, when many photoreceptor progenitors have normally stopped proliferating and have fully differentiated, *nfkb1* expression is still higher in *miR-18a^mi5012^* retinas (WT 1.03 ± 0.07 SD, *miR-18a^mi5012^* 1.33 ± 0.22 SD, p=0.044, n=3) (Figure 9d). Taken together, these results indicate that following photoreceptor injury, in *miR-18a^mi5012^* retinas compared with WT, there is both a higher level of inflammatory pathway activity and a prolonged inflammatory response.

**Fig. 9.**
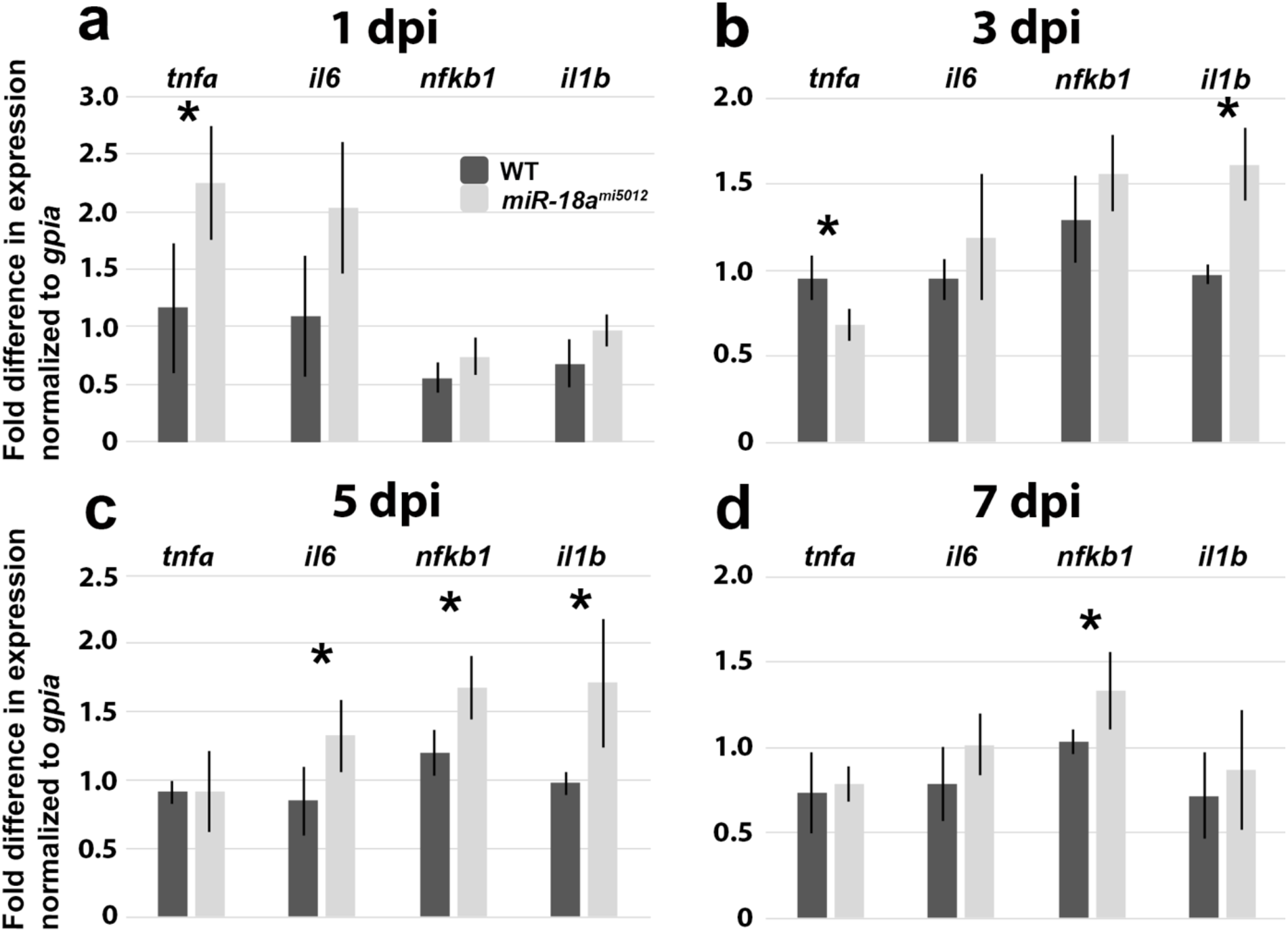
RT-qPCR showing expression of inflammatory molecules in WT and *miR-18a^mi5012^* retinas at 1, 3, 5 and 7 dpi. (a-d) RT-qPCR showing fold differences in expression of the inflammatory genes *tnfα* (*tnfa*), *il6*, *nfkb1* and *il1β* (*il1b*) compared with WT 1dpi, normalized to the housekeeping gene *gpia*, in WT and *miR-18a^mi5012^* retinas at 1 (a), 3 (b), 5 (c) and 7 dpi (d). Numbers and p values for significant differences are show in the text. Error bars represent standard deviation and asterisks indicate significant differences (Student’s t-test, p<0.05)

### Loss of *miR-18a* results in a prolonged microglia/macrophage response

Microglia respond to retinal injury by undergoing a morphological change from ramified to amoeboid, migrating to the injury site and phagocytosing apoptotic cells, and releasing pro-and anti-inflammatory cytokines and other signaling molecules [reviewed in 24]. The prolonged inflammation in *miR-18a^mi5012^* retinas during photoreceptor regeneration, and the central role that microglia play in the injury-induced inflammatory response, suggest that *miR-18a* might regulate the microglia response to injury. To determine if *miR-18a* is expressed in microglia/macrophages following photoreceptor injury, in situ hybridization for *miR-18a* was used in combination with immunolabeling with the 4c4 antibody that labels microglia/macrophages in zebrafish. At 3 dpi, when microglia are active and *miR-18a* expression is the highest, *mir-18a* is expressed rarely in microglia/macrophages (arrowhead, Figure 10a), suggesting that *miR-18a* does not directly regulate microglia. Next, to determine if loss of *miR-18a* alters the response of microglia/macrophages to photoreceptor injury, these cells were immunolabeled and counted in WT and *miR-18a^mi5012^* retinas at 3, 5, 7, 10 and 14 dpi. The results showed no differences in the total numbers of microglia/macrophages at 3, 5 or 7 dpi (Figure 10b-g). In contrast, at 10 dpi, *miR-18a^mi5012^* retinas had more microglia/macrophages (Figure 10h, i) than WT, and this difference was even more pronounced at 14 dpi (Figure 10j, k). Additionally, in *miR-18a^mi5012^* retinas compared with WT, there were more microglia/macrophges in the outer retina (RPE + ONL) at 7dpi, in the inner retina (INL + GCL) at 10 dpi, and in both the outer and inner retinas at 14 dpi (Figure 10f-k). These data indicate that *miR-18a* does not regulate the early response of microglia/macrophages to photoreceptor injury, but that loss of *miR-18a* results in prolonged activity of microglia/macrophages.

**Fig. 10.**
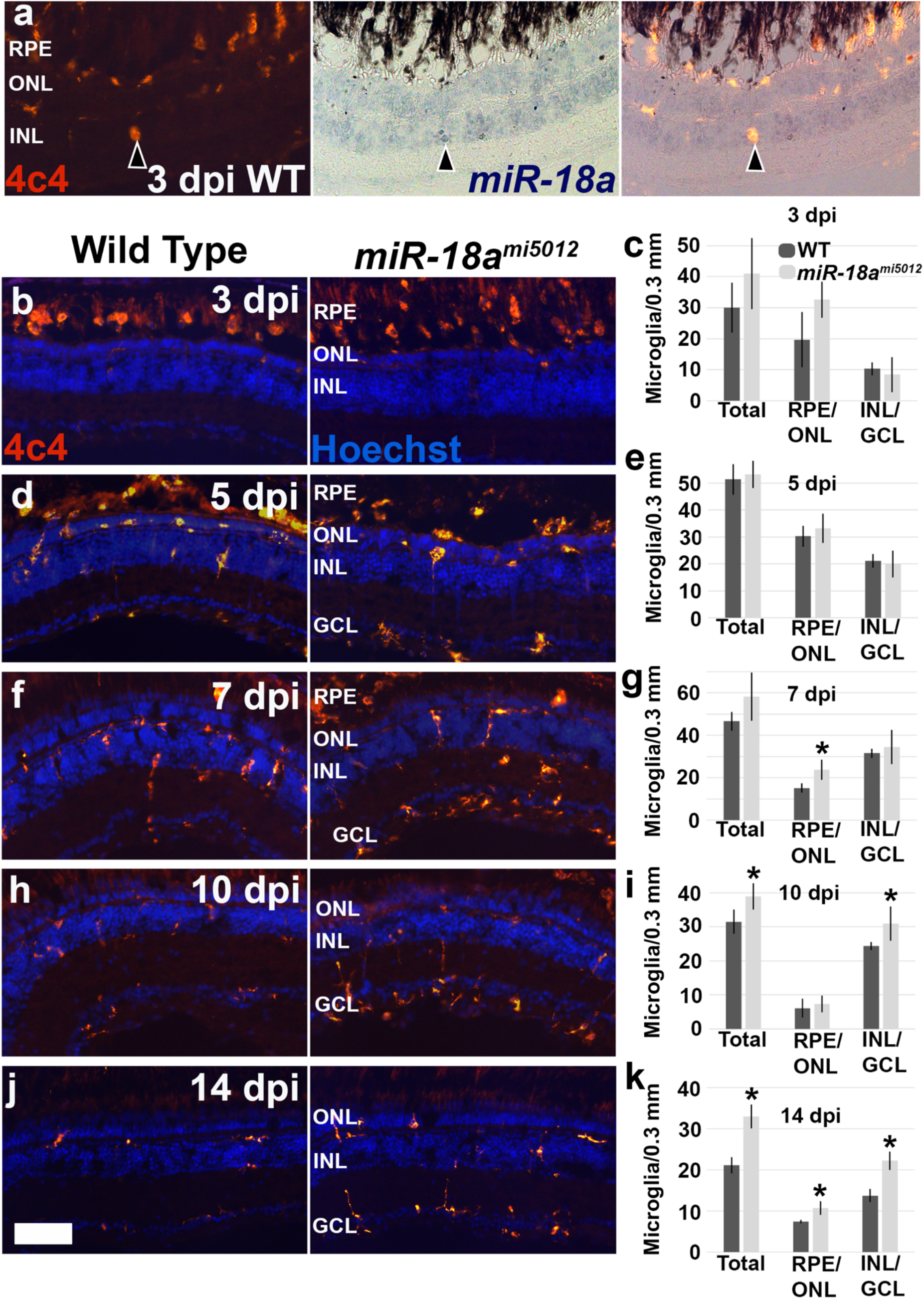
Expression of *miR-18a* in microglia, and comparison of the microglia response to photoreceptor injury in WT and *miR-18a^mi5012^* retinas. (a) In situ hybridization for miR-18a and co-localization with 4C4 labeling in microglia (red) in only one microglial cell in this image (arrowhead). (b-k) Microglia labeling with 4C4 antibody (red) in WT and *miR-18a^mi5012^* retinas at 3 dpi (b, c) (total microglia WT 30.0 ± 8.0 SD, *miR-18a^mi5012^* 41.0 ± 11.4 SD cells/0.3 mm, p=0.12, n=3), 5 dpi (d, e) (total microglia WT 51.3 ± 5.7 SD, *miR-18a^mi5012^* 53.2 ± 5.0 SD cells/0.3 mm, p=0.69, n=3), 7 dpi (f, g) (total microglia WT 46.6 ± 4.4 SD, *miR-18a^mi5012^* 58.1 ± 11.3 SD cells/0.3 mm, p=0.17, n=3), 10 dpi (h, i) (total microglia WT 31.5 ± 3.5 SD, *miR-18a^mi5012^* 38.9 ± 3.9 SD cells/0.3 mm, p=0.03, n=3), and 14 dpi (total microglia WT 21.2 ± 2.0 SD, *miR-18a^mi5012^* 33.0 ± 2.9 SD cells/0.3 mm, p=0.001, n=4). Additionally, microglia were counted differentially in the outer retina (RPE/ONL) and the inner retina (INL/GCL) and significant differences are as follows: 7dpi RPE/ONL (WT 15.1 ± 2.2 SD, *miR-18a^mi5012^* 23.7 ± 4.7 SD cells/0.3 mm, p=0.047, n=3), 10 dpi INL/GCL (WT 24.3 ± 1.2 SD, *miR-18a^mi5012^* 30.9 ± 5.0 SD cells/0.3 mm, p=0.046, n=3), 14 dpi (RPE/ONL—WT 7.4 ± 0.4 SD, *miR-18a^mi5012^* 10.8 ± 1.6 SD cells/0.3 mm, p=0.008, n=4; INL/GCL—WT 13.8 ± 1.6 SD, *miR-18a^mi5012^* 22.3 ± 2.2 SD cells/0.3 mm, p<0.001, n=4). Cells were counted in 0.3 mm of linear retina at each time point; Error bars represent standard deviation and asterisks indicate significant differences (Student’s t-test, p<0.05). RPE—retinal pigmented epithelium, ONL—outer nuclear layer, INL—inner nuclear layer, GCL—ganglion cell layer; *scale bar:* 50 mm

### Supressing inflammation in *miR-18a* mutants rescues both the excess proliferation and delayed photoreceptor regeneration

The increased and prolonged expression of inflammatory genes and prolonged microglia/macrophage response in the injured *miR-18a^mi5012^* retina led to the hypothesis that, in the absence of *miR-18a*, prolonged and excessive inflammation is causally related to the excess proliferation and delayed photoreceptor maturation observed in *miR-18a* mutants. To test this hypothesis, dexamethasone was used to suppress inflammation in WT and *miR-18a^mi5012^* fish from 2 to 6 dpi, the interval during which *miR-18a* expression is upregulated in injured retinas (see Figure 1a, b) and when in *miR-18a^mi5012^* retinas inflammatory genes are expressed higher levels than in WT retinas (see Figure 9). Fish were then exposed to 5 mM BrdU from 6 to 7 dpi, to label proliferating cells, and *in situ* hybridization was used to label mature rod or cone photoreceptors. Compared with controls, the dexamethasone treatment fully rescued the excess proliferation in *miR-18a^mi5012^* retinas, reducing the number of BrdU-labeled cells to that observed in controls (Figure 11a, b). Dexamethasone treatment also reduced the number of BrdU-labeled cells in WT retinas by a smaller amount (Figure 11a, b). Further, dexamethasone treatment rescued the delay in cone maturation and regeneration at 7 dpi, matching the number of ZPR-1-labeled mature cones with WT control levels (Figure 11c, e). Dexamethasone treatment had no effect on the number of cone cells in WT retinas (Figure 11c, e) and had no effect on the numbers of Zpr-3-labeled rod cells (Figure 11d). Together, these data show that in the injured retina, the *miR-18a* mutant phenotype is fully rescued through inflammatory suppression with dexamethasone, indicating that *miR-18a* function is mediated through inflammatory regulation.

**Fig. 11.**
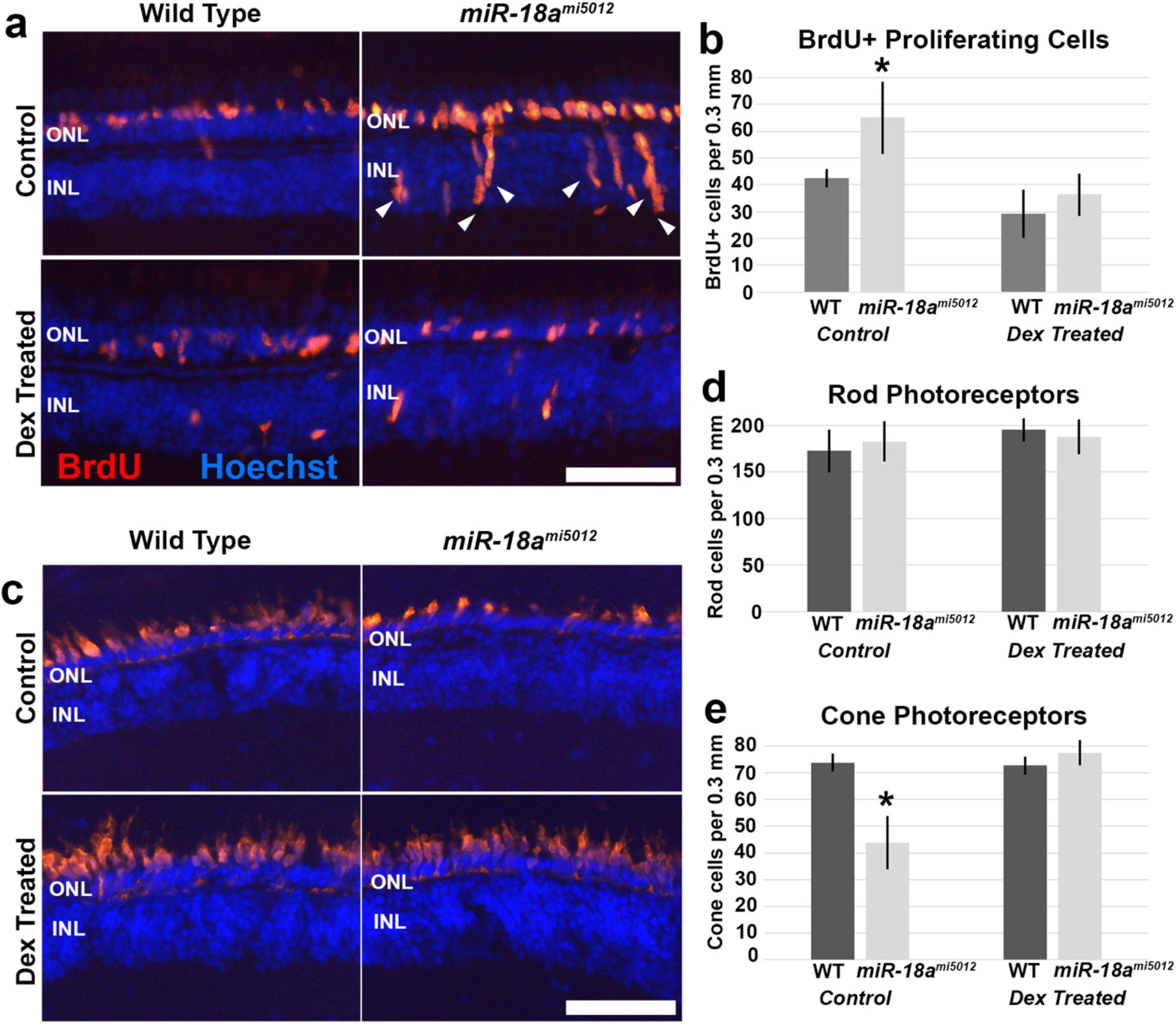
Rescue of the 7 dpi *miR-18a^mi5012^* phenotype by inflammatory suppression with dexamethasone. (a) BrdU immunolabeling (orange-red) in retinal cross sections of WT and *miR-18a^mi5012^* fish exposed to BrdU from 6-7 dpi in control fish (top panels) and dexamethasone treated (Dex) fish (treated from 2-6 dpi to suppress inflammation) (bottom panels); arrowheads show BrdU-labeled cells in the INL of *miR-18a^mi5012^* retinas (to right) that are largely absent from WT and Dex treated retinas. (b) Quantification of BrdU-labeled cells in retinal cross sections of WT and *miR-18a^mi5012^* fish exposed to BrdU from 6-7 dpi in control and dexamethasone treated fish (WT control 42.5 ± 3.5 cells/0.3 mm vs. *miR-18a^mi5012^* control 65.0 ± 13.5 cells/0.3 mm, p=0.009, n=4; WT control 42.5 ± 3.5 cells/0.3 mm vs. *miR-18a^mi5012^* dexamethasone treated 36.2 ± 7.7 cells/0.3 mm, p=0.096, n=4); (WT control 42.5 ± 3.5 cells/0.3 mm vs. WT dexamethasone treated 29.2 ± 9.0 cells/0.3 mm, p=0.020, n=4). Cells were counted over 0.3 mm linear retina; error bars represent standard deviation and asterisks indicate significant differences (Student’s t-test, p<0.05). (c) Immunolabeling for mature cones (ZPR-1/Arr3a) in retinal cross sections of WT and *miR-18a^mi5012^* control fish (top panels) and dexamethasone treated fish (bottom panels). (d) Quantification of mature rods (ZPR-3/Rho-immunolabeled) showing no significant differences across groups or treatments. (e) Quantification of mature cones (ZPR-1/Arr3a-immunolabeled) in retinal cross sections of WT and *miR-18a^mi5012^* control and dexamethasone treated fish (WT control 73.8 ± 3.4 cells/0.3 mm vs. *miR-18a^mi5012^* control 43.9 ± 10.0 cells/0.3 mm, p=0.001, n=4; WT control 73.8 ± 3.4 cells/0.3 mm vs. *miR-18a^mi5012^* dexamethasone treated 77.6 ± 4.8 cells/0.3 mm, p=0.254, n=4); (WT control 73.8 ± 3.4 cells/0.3 mm vs. WT dexamethasone treated 72.7 ± 3.3 cells/0.3 mm, p=0.670, n=4). Cells were counted over 0.3 mm linear retina; error bars represent standard deviation and asterisks indicate significant differences (Student’s t-test, p<0.05). Abbreviations: RPE—retinal pigmented epithelium, ONL—outer nuclear layer, INL—inner nuclear layer; *scale bars:* 50 mm

## DISCUSSION

Recent studies have dramatically improved our understanding of the mechanisms that govern the Müller glia-based neuronal regeneration in the injured zebrafish retina [reviewed in 4,43,10]. These results led directly to studies showing that mammalian Müller glia possess latent regenerative potential that can be augmented by forced reprogramming, and that MG-derived progenitors can regenerate some neurons (including photoreceptors) [7,25,8,44–46], though this neuronal regeneration is very inefficient [9, 43]. In the zebrafish retina, inflammation is required for neuronal regeneration [47,17,18,48,20,19,23,21,22], but when inflammation is mis-regulated in the zebrafish retina, this leads to aberrant proliferation of MG-derived progenitors and altered photoreceptor regeneration [21, 20], showing the importance of precise inflammatory regulation in controlling these events.

Several miRNAs have been identified as important regulators of inflammation and could be key neuroinflammatory regulators in the injured retina [26, 27], but there are few studies linking miRNAs with retinal inflammation. *miR-18a* is predicted to interact with mRNAs of more than 25 inflammation-related molecules (http://www.targetscan.org/fish_62), suggesting that *miR-18a* could regulate neuroinflammation. However, the roles of *miR-18a* in injury-induced inflammation and photoreceptor regeneration had not been previously investigated.

The objective of the current study was to determine the role of *miR-18a* in photoreceptor regeneration and to determine if it regulates inflammation during this response. Following photoreceptor injury, *miR-18a* expression increases throughout the retina, and is expressed in the INL and ONL, in Müller glia as early as 1 dpi that divide around this time, and in both Müller glia and proliferating MG-derived progenitors at 3-5 dpi, indicating that *miR-18a* functions during key times of cell division. RT-qPCR showed increased expression of both the *miR-18a* precursor (*pre-miR-18a*) and mature *miR-18a* at 3 and 5 dpi, showing that, as in the developing retina [30], the precursor and mature miRNA molecules are expressed proportionally. Previously, using RNA-deep sequencing (RNA-seq), Rajaram et al. [11] also showed a modest increase in *miR-18a* expression by 3 dpi, but the photolytic lesioning methods and timing were different than in the current study, and RT-qPCR (used here) is considered to be more specific and sensitive than RNA-seq. Uninjured WT and *miR-18a^mi5012^* adult retinas have identical numbers of mature photoreceptors and, by 14 days after photoreceptor injury, WT and *miR-18a^mi5012^* retinas regenerate equivalent numbers of photoreceptors. This shows that during photoreceptor regeneration, as in the developing embryonic retina [30], *miR-18a* does not regulate the overall number of photoreceptors that differentiate. In the injured adult retina the loss of *miR-18a* results in increased proliferation among MG-derived progenitors and delayed photoreceptor differentiation. These differences indicate that in the adult retina, *miR-18a* functions through pathways that are specific to the injury response.

In *miR-18a^mi5012^* retinas compared with WT, the number of dividing progenitors is identical at 3 dpi, the peak of the proliferative response but, in *miR-18a^mi5012^* and *miR-18a* MO-injected retinas compared with WT/controls, more of these cells continue to proliferate at 7 and 10 dpi, when most photoreceptor progenitors have normally exited the cell cycle. These data indicate that *miR-18a* regulates proliferation of MG-derived progenitors and that loss of *miR-18a* results in a prolonged proliferative response. This function of *miR-18a* to limit progenitor proliferation differs from the function of *miR-18a* in some cancers, in which *mirR-18a* promotes the proliferation of tumor cells [49, 50] including glioblastoma cells [51]. Also, in the developing mouse neocortex, the *miR-17-92* cluster, which includes *miR-18a*, has been shown to promote proliferation of neuronal progenitors [52]. Interestingly, however, *miR-18a* has also been shown to suppress cell proliferation in pancreatic progenitor cells [49] and myoblast cells [53], and it has anti-proliferation/anti-tumor effects in colorectal and breast cancers [54,50,55]. The effects of *miR-18a* on cell proliferation are, therefore, dependent on the cell type and injury/disease state of the tissue involved.

In *miR-18a^mi5012^* retinas, the prolonged proliferation among MG-derived progenitors results in a delay in photoreceptor regeneration and maturation, and an excess number of MG-derived progenitors. A significant number of these progenitors remain in the INL, consistent with data showing that Müller glia derived progenitors are multipotent and can generate multiple cell types even though they preferentially generate the neurons that were ablated [56–58]. Chemokines, released in response to cell death, guide the migration of neural progenitors [59]. While it is unknown if *miR-18a* regulates chemokines directly, an altered chemokine milieu in retinas of the *miR-18a* mutants could lead to aberrant progenitor migration, resulting in more progenitors failing to migrate to the ONL.

The excess proliferation of progenitors and higher expression of key inflammatory genes in *miR-18a^mi5012^* retinas is similar to what was observed in *mmp9* mutants, but there are also some important differences. First, the excess progenitors in *miR-18a^mi5012^* retinas do not generate excess photoreceptors, and this differs from what is observed in *mmp9* mutant retinas [21]. This difference indicates that *miR-18a* and Mmp9 may also differentially regulate molecules downstream of pathways that regulate inflammation and cell proliferation. In the subventricular zone of the CNS, Mmp9 has been found to regulate differentiation of neuronal progenitor cells [60], indicating that it has some capacity to regulate neuronal differentiation. As a microRNA, *miR-18a* likely regulates several classes of molecules involved in the various steps of neuronal regeneration, and members of the *miR-17-92* cluster of miRNAs, which includes *miR-18a*, are key regulators of neurogenesis, involved in cell proliferation, progenitor fate determination and differentiation [reviewed in 61]. Previous work found that NeuroD governs the cell cycle and differentiation among photoreceptor progenitors during both embryonic development and regeneration [62, 42] and that *miR-18a* regulates NeuroD protein levels in the embryonic retina [30]. Downstream of inflammatory pathways, *miR-18a* is also likely to regulate NeuroD in the injured retina, and this could affect differentiation of photoreceptors and other cells. The dominant phenotype in injured adult *miR-18a^mi5012^* retinas, however, is the increased inflammation and cell proliferation. Pathways downstream of inflammation have not yet been investigated.

A second key difference between the *miR-18a^mi5012^* and *mmp9* mutant retinal phenotypes is that, although both mutations result in an elevated inflammatory response following photoreceptor injury, they result in higher expression of different inflammatory molecules and at different time points. In *miR-18a^mi5012^* retinas, *tnfα* expression is increased only at 1 dpi, differing markedly from observations in *mmp9* mutant retinas that have increased expression of *tnfα* (but not other inflammatory molecules) at all post-injury time points [21]. Also in contrast to *mmp9* mutants, *miR-18a^mi5012^* retinas have increased expression of other cytokines (*il6* and *il1b*) and the cytokine regulator *nfkb1* at 5 dpi, a time point when inflammation is typically subsiding and progenitors are beginning to differentiate. Further, *nfkb1* expression continues to be higher in *miR-18a^mi5012^* retinas at 7dpi, when widespread differentiation is typically occurring among photoreceptor progenitors. These data indicate that, like Mmp9, *miR-18a* negatively regulates inflammation, leading to a converging phenotype (excess neuronal progenitors). However, *miR-18a* regulates different inflammatory molecules than Mmp9 and, in *miR-18a^mi5012^* retinas, inflammation is prolonged.

In situ hybridation showed that at 3 dpi, *miR-18a* is not expressed in most retinal microglia/macrophages but that, in *miR-18a^mi5012^* fish compared with WT, the microglia/macrophage response to retinal injury is prolonged and there are differences in the localization of microglia/macrophages. The lack of *miR-18a* expression in microglia/macrophages indicates that *miR-18a* likely does not regulate microglia directly, but the presence of more [total] microglia/macrophages in *miR-18a^mi5012^* retinas than WT at 10 and 14 dpi, the presence of more microglia/macrophages in the outer retina at 7 dpi, the inner retina at 10 dpi, and throughout the retina at 14 dpi, and the increased expression of inflammatory cytokines at 1, 3, 5 and 7 dpi, suggest that *miR-18a* indirectly regulates the microglia response to photoreceptor injury. This indirect regulation of microglia could be mediated through Müller glia, RPE and/or dying photoreceptors, all of which have the ability to regulate microglia activity via chemokine secretion [63–65]. The differences observed in inflammatory cytokine expression between *miR-18a^mi5012^* and WT retinas likely results from differences in the extent and duration of the microglia response, regulated indirectly through *miR-18a* and other cell types. An example of such a regulatory mechanism could be through the chemokine CXCL12, a predicted regulatory target of *miR-18a* (http://www.targetscan.org/fish_62) that has been shown to be expressed in radial glia [66] and RPE cells [67], and mediates microglia activity and migration through the CXCR4 receptor. CXCR4 is also a predicted regulatory target of *miR-18a* (http://www.targetscan.org/fish_62), making the CXCL12/CXCR4 axis a strong candidate mechanism through which *miR-18a* could regulate microglia activity and inflammation.

The prolonged expression of important inflammatory molecules in *miR-18a^mi5012^* retinas during photoreceptor regeneration indicates that *miR-18a* functions during the resolution phase of the inflammatory response, potentially regulating a negative feedback loop that resolves inflammation and restores homeostasis. Resolving neuroinflammation promotes normal tissue repair and if acute inflammation remains unresolved, it can result in inadequate repair or further neuronal damage [68, 24]. Following photoreceptor injury, expression levels of *nfκb1* and certain cytokines are normally upregulated and peak between 24 and 48 hpi and then their expression levels decrease, returning to homeostasis sometime after 7 dpi [see 21]. The removal of NFκB activity and inflammatory cytokine signaling are critical steps in resolving inflammation [69], indicating that the time frame between 48 hpi and 7 dpi in the retina is key for regulating this process. In *miR-18a^mi5012^* retinas, the higher expression of *nfκb1* and inflammatory cytokines at 5 dpi compared with WT, and the continued higher expression of *nfκb1* at 7 dpi, indicate that resolution of the inflammatory response is delayed in the absence of *miR-18a*. Treatment of *miR-18a^mi5012^* fish with dexamethasone was a means to chemically resolve retinal inflammation during photoreceptor regeneration, and this fully rescued the excess proliferation and delayed photoreceptor regeneration in the *miR-18a^mi5012^* retina. This indicates that *miR-18a* functions during the resolution phase and plays a role in regulating key aspects of inflammation.

Inflammation in the retina following photoreceptor death is generated by the release of cytokines from microglia [70], Müller glia [71], retinal pigmented epithelium (RPE) [reviewed in 72] and dying cells [17]. An underlying assumption is that these cells are responsible for the elevated and prolonged inflammation in the *miR-18a^mi5012^* retinas, however, given that *miR-18a* expression is elevated relatively ubiquitously in the injured retina, the increased and prolonged inflammation in the mutants could originate from other cellular types. Müller glia are known to secrete inflammatory cytokines in response to retinal injury [23,17,19] and cytokine signals from microglia and/or dying cells are necessary for Müller glia to divide [17, 19]. The initial expression of *miR-18a* in Müller glia and other retinal cells might, therefore, be a mechanism to limit the level of retinal inflammation by regulating the secretion and responses to certain inflammatory cytokines. The highest expression of *miR-18a* occurs between 3 and 5 dpi, during the resolution phase of retinal inflammation [see 21]. Based on its 7-base seed sequence (CACCUUA), *miR-18a* is predicted to interact directly with mRNAs of more than 25 molecules that function in inflammatory pathways (http://www.targetscan.org/fish_62), some of which are cytokines and other intercellular signaling molecules (e.g. *1l-16*, *cxcl12a, bmp6*), but many of which are transmembrane receptors or molecules that function downstream of inflammatory cytokines (e.g. *Il-7r*, *cxcr4a*, *tnfaip3*). It is therefore probable that *miR-18a* not only reduces inflammatory signals from Müller glia and/or other cells (e.g. microglia, dying photoreceptors), but also reduces the effects of those inflammatory signals on the MG-derived progenitors.

The role of *miR-18a* in regulating progenitor proliferation, photoreceptor regeneration and inflammation in the injured retina adds to the growing body of work showing that several miRNAs are key inflammatory regulators and/or play critical roles in neuronal regeneration. The miRNA *let-7* was one of the first identified to regulate the response of Müller glia to retinal injury in zebrafish. In the retina, *let-7* functions by suppressing dedifferentiation and division of Müller glia until its levels are reduced by Lin-28 to initiate the regeneration response [14]. Similarly, following retinal injury, *miR-216a* prevents Müller glia reprogramming and proliferation, and reducing *miR-216a* is necessary and sufficient to activate Wnt signaling and initiate MG division [73]. After MG initially divide, *miR-203* was then found to suppress proliferation among MG-derived progenitors, and its level must be reduced to allow adequate cell proliferation and neuronal regeneration [12]. These studies show that several miRNAs function to prevent a premature or aberrant regeneration response and to limit cell proliferation. In line with this, the current study shows that *miR-18a* limits proliferation among MG-derived progenitor cells, but appears to function later than the previously studied miRNAs to limit the duration of the proliferative response rather than prevent it from initially occurring. This is likely accomplished by *miR-18a* functioning during the resolution phase of inflammation and limiting the duration of the inflammatory response. While *miR-18a* is the first known miRNA to play important roles in both neuronal regeneration and inflammation in the injured retina, several other miRNAs are known to be key regulators of inflammation [74] and could be important for future retinal studies. For example, in patients with diabetic retinopathy, decreased *miR-146b-3p* was associated with increased vascular inflammation [75] and in the healthy and degenerating mouse retina, *miR-223* is an important inflammatory suppressor [76]. MicroRNAs can also have pro-inflammatory functions; an example is *miR-155*, which was shown to promote inflammation in the degenerating mouse retina, Conversely, inhibiting *miR-155* in this context was neuroprotective [77]. In addition to regulating inflammation, miRNA profiling studies combined with broad blocking of miRNA function (e.g. Dicer knockout) are revealing the potential importance of many miRNAs in regulating neuronal regeneration in the zebrafish retina [11] and in reactive gliosis and understanding the mechanisms that prevent Müller glia reprogramming and neuronal regeneration in the mammalian retina [78]. Studies like these are bringing miRNA research to the forefront and showing the complexities of understanding pathways that govern neuronal regeneration.

In conclusion, this study is the first to show that the presence of a single miRNA is critical for inflammatory resolution in the injured/regenerating retina and is the first to show that *miR-18a* plays important roles in both inflammation and photoreceptor regeneration. This work adds to the growing body of knowledge that miRNAs are key regulators of neurogenesis throughout the central nervous system [79] and neuronal regeneration in the retina [15]. Importantly, the differential responses of Müller glia to inflammation following retinal injury in zebrafish compared with mammals could be key to unlocking the potential for mammalian Müller glia to robustly regenerate neurons. Like *miR-18a*, other miRNAs could be potent inflammatory regulators in the injured retina and may be key to fully augmenting the regenerative potential of the mammalian retina.

## DECLARATIONS

### Ethics approval

All experimental procedures were approved by the University of West Florida Institutional Animal Care and Use Committee.

### Consent to Participate

N/A

### Consent for publication

N/A

### Informed Consent

N/A

### Availability of data and materials

All data and *miR-18a^mi5012^* fish are freely available upon request

## Funding

This research was supported by the following grants: NIH 1R15EY031089-01 (SMT), NIH T32EY013934 (SMT); NIH F30EY031142 (ACK), NIH P30EY004068 (RT), NIH R21 EY031526 (RT), NIH P30EYO7003 (PFH), NIH R01EY07060 (PFH), and an unrestricted grant from the Research to Prevent Blindness, New York (RT and PFH).

### Authors’ Contributions

All authors contributed to the study conception and design. Material preparation, data collection and analysis were performed by Evin Magner, Pamela Sandoval-Sanchez, Ashley Kramer, Ryan Thummel and Scott M. Taylor. The first draft of the manuscript was written by Evin Magner, Pamela Sandoval-Sanchez, and Scott M. Taylor, and all authors commented on previous versions of the manuscript. All authors read and approved the final manuscript.

## Acknowledgements

The authors would like to thank James Hammond at UWF for facilities support. We also thank Karen Gibbs, the RAE Office and the Hal Marcus College of Science and Engineering for facilitating administrative, technical and financial support.

---

**Fig. S1.**
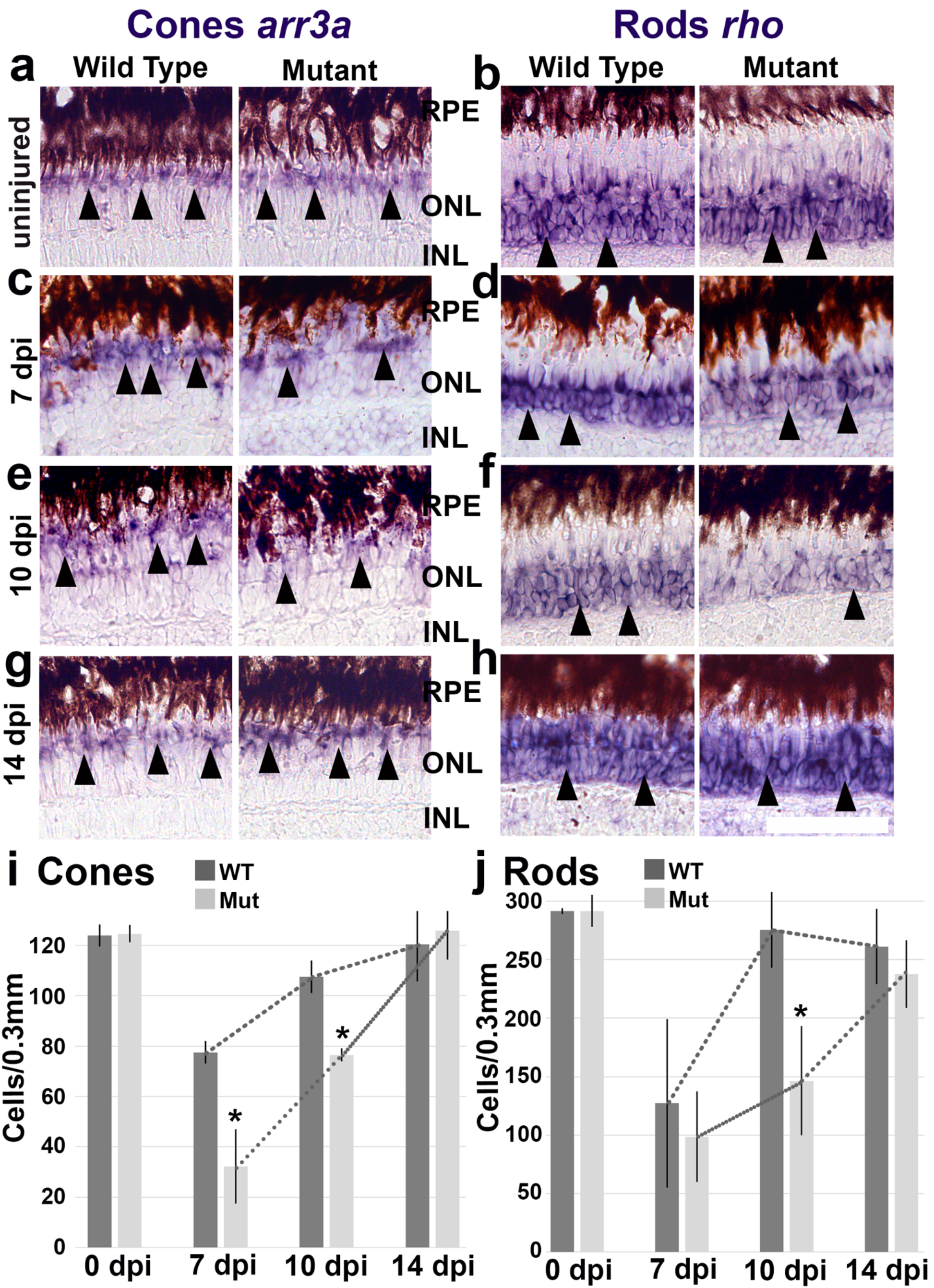
*In situ* hybridizations and quantification of mature cone and rod photoreceptors in WT and *miR-18a^mi5012^* retinas in uninjured retinas and at different numbers of days post-injury (dpi). (a-h) *In situ* hybridizations and (i, j) quantification for mature cones (*arr3a*) and rods (*rho*) in WT and *miR-18a^mi5012^* retinas in uninjured reinas (a, b, i, j) (cones—WT 123.8 ± 4.3 SD, *miR-18a^mi5012^* 124.6 ± 3.4 SD cells/0.3 mm, p=0.81, n=3; rods—WT 291.7 ± 2.3 SD, *miR-18a^mi5012^* 291.8 ± 13.8 SD cells/0.3 mm, p=0.99, n=3), at 7 dpi (c, d, i, j) (cones—WT 77.5 ± 4.5 SD, *miR-18a^mi5012^* 32.3 ± 14.7 SD cells/0.3 mm, p=0.001, n=3; rods—WT 127.1 ± 72.1 SD, *miR-18a^mi5012^* 98.5 ± 38.6 SD cells/0.3 mm, p=0.51, n=3), at 10 dpi (e, f, i, j) (cones—WT 107.6 ± 6.5 SD, *miR-18a^mi5012^* 76.4 ± 2.5 SD cells/0.3 mm, p=0.001; rods—WT 275.6 ± 32.5 SD, *miR-18a^mi5012^* 146.4 ± 46.7 SD cells/0.3 mm, p=0.017, n=3) and at 14 dpi (g-j) (cones—WT 120.3 ± 14.7 SD, *miR-18a^mi5012^* 125.7 ± 11.4 SD cells/0.3 mm, p=0.462; rods—WT 261.3 ± 32.1 SD, *miR-18a^mi5012^* 237.7 ± 28.8 SD cells/0.3 mm, p=0.241, n=3). Arrowheads show examples of labeled photoreceptors. Photoreceptor counts in retinal cross sections in the center of the lesioned area (cells per 0.3 mm of linear retina). Error bars represent standard deviation and asterisks indicate significant differences (Student’s t-test, p<0.05). Dotted lines on the graph connect the tops of the bars to show the trends. Abbreviations: RPE—retinal pigmented epithelium, ONL—outer nuclear layer, INL—inner nuclear layer; *scale bar:* 50 μm

**Fig. S2.**
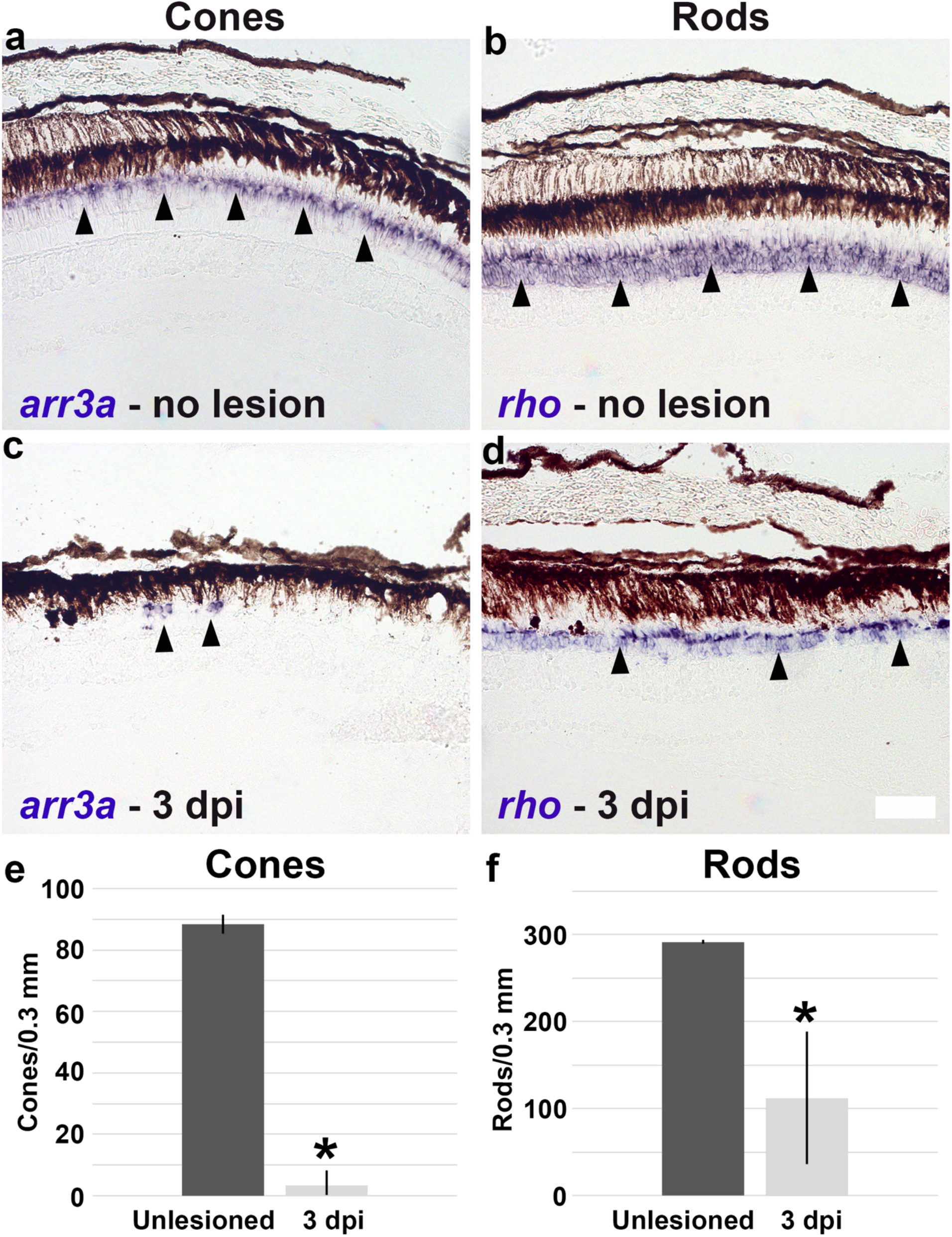
Efficiency of photolytic lesioning of photoreceptors in WT fish. (a, b) In situ hybridization for *arr3a* (cones) and *rho* (rods) in unlesioned retinas. (c, d) In situ hybridization for *arr3a* (cones) and *rho* (rods) in lesioned retinas at 3 dpi. (e, f) Quantification of mature cones (*arr3a*-labeled) and rods (*rho*-labeled) in retinal cross sections in unlesioned retinas and at 3 dpi (cones—unlesioned 88.4 ± 3.1 SD, 3 dpi 3.2 ± 5.0 SD cells/0.3 mm, p<0.0001, n=3; rods—unlesioned 291.7 ± 2.3 SD, 3 dpi 112.1 ± 76.5 SD cells/0.3 mm, p<0.015, n=3). Cells were counted over 0.3 mm linear retina; error bars represent standard deviation and asterisks indicate significant differences (Student’s t-test, p<0.05); *scale bar:* 50 μm

